# Dichotomy in TCR V-domain dynamics binding the opposed inclined planes of pMHC-II and pMHC-I α-helices

**DOI:** 10.1101/2023.01.16.524284

**Authors:** Joseph S. Murray

**Author notes:** Address: 10880 Willshire Blvd., Suite 1101, Los Angeles, CA 90024.

## Abstract

Ligand recognition by the human α/β T-cell antigen receptor (TCR) heterodimer protein, unlike the surface immunoglobulin (sIg) B-cell receptor, is not governed by relative binding affinity. Its interaction with the peptide (p) plus major histocompatibility complex (MHC) protein (abbrev. pMHC) likely involves some different molecular mechanism linking pMHC binding to T-cell functions. Recent analytical geometry of TCR:pMHC-II solved crystallographic structures (*n* = 40) revealed that each variable (V)-domain is bound in similar, yet mathematically unique orientations to its target pMHC groove. The relative position of the central cysteine of each V-domain was examined by multivariable calculus in spherical coordinates, where a simple volume element (*dV*) was found to describe clonotypic geometry with pMHC-II. Here, the study was expanded to include TCR:pMHC-I structures, and to model a physical mechanism, specifically involving the two directionally opposed *inclined planes* (IP) manifest by the two major α-helices prominent in both MHC-I and MHC-II proteins. Calculations for rotational torque of each V-domain, together with acceleration up and down the slopes of both MHC α-helices were used to estimate the time a given V-domain spends sliding down its cognate MHC IP. This V-domain rotation/sliding mechanism appears to be quantitatively unique for each TCR:pMHC V-domain (*n* = 40). However, there is an apparent and common dichotomy between the mobility of each V-domain with respect to the two classes of MHC proteins. Evolutionary motifs in the MHC helices support that the V-domains negotiate the opposed inclined planes of pMHC ligands in clonotypic fashion. Thus, this model is useful in understanding how mechanical forces are linked to TCR function.

## Introduction

The capacity to recognize subtle structural differences between pathogens and to ‘recall’ them, such that an effective host response can kill and clear pathogens, is the providence of the ‘adaptive’ immune system (ref. 1). The word *adaptive* derives from the property of the system to “re-configure” itself in *real time*. In other words, this system of complex proteins, and the cellular paradigms upon which they function, appears to *adapt* over time (within the individual) to optimize survival against a world of rapidly reproducing (rapidly evolving) pathogenic microorganism. This capacity is highly unusual compared to most genes [1–7].

*Adaptive immunity* is based on the biology that precursor T cells and B cells go through a process of development wherein their antigen receptors are “individualized” within each clone. This is essential. Without this process of somatic DNA rearrangement in the nucleus of every developing clone of T cell and B cell, the immune system would not have the enormous structural diversity in the variable domains of these two types of special proteins required for the job of specific pathogen recognition. In essence, the adaptive immune system is ‘hard wired’ for *clonal selection*, first in the thymus or bone-marrow, then later, *by the antigen* for the best receptors to use against it [1–4]. This might seem counter-intuitive from the pathogen’s point of view, but pathogens (e.g., HIV-1) continuously evolve their peptide components to precisely avoid this *adaptive* capacity of the vertebrate immune system [8]. In a way, it is a ‘steady state’ in the natural history of host and pathogen, where the overall system has reached mechanisms that allow both to survive [9]. Unfortunately, even with an adaptive immune system, there is high morbidity and mortality, particularly with ‘emerging’ pathogens for which few individuals have so adapted clonal populations adequate to fight otherwise transient pathogenicity. This is reflected in the need for vaccination, which artificially adapts the host system, such that upon infection the pathogen is met with a sufficiently prepared repertoire of TCR and sIg capable of killing and eradication of the pathogen for which the vaccine was synthesized [1, 10].

Clonotypic rotational volumetric density (*dV*) of both V-domains within a panel (*n* = 40) of human α/β T-cell antigen receptors in protein-to-protein complex with their ‘target’ pMHC class II ligands was recently reported [11]. Earlier, expression quantitative trait locus (eQTL) mapping showed that TCR V-gene segment usage is linked (*in trans*) to the extremely polymorphic MHC (both MHC-II and (less so) MHC-I) antigen-presenting genes [12]. There is evidence that both the TCR and the larger ‘TCR-dimer:CD3-dimers’ complex are influenced by *mechanical* forces operating at the junction between opposing T-cell and antigen presenting cell (APC) biological membranes [13–17]. The CD3 dimers have tyrosine phosphorylation sites on long cytoplasmic tails (ITAM) motifs that are themselves subject to conformational and combinatorial mechanisms known to function in recognition by CD4/CD8 associating tyrosine kinases (Lck), phosphatases (CD45), which are further linked to cascades (transduction pathways) delivered by CD28 (PIP3), Ca++ flux (CaM), further phosphorylation and dephosphorylation (PKC, LAT, CD6), combinatorial kinase activation (ZAP70, Erk), and thought to lead to the activation of specific promotors (*via* NF-κB, Jun, NFAT) for genes involved in the basic mechanisms of T-cell effector functions, e.g., induction of clonal division (proliferation), the activation of direct infected-cell killing by T cells, and genes for major cell to cell communications, cytokines [1, 17, 18]. Crucially, these down-stream mechanisms are all theoretically controlled by the initial interaction of the α/β component of the TCR with the pMHC-I or pMHC-II ligand, whose antigen-specific recognition by a given T-cell clone is ultimately a result of the somatic DNA rearrangements encoding the TCRα and TCRβ protein chains. Rearrangements occur in the foetal thymus prior to negative and positive selection by self-pMHC ligands on bone-marrow derived and epithelial cells, which select only those TCR-bearing clones that recognize the host’s particular MHC genotype [1, 2]. This mechanism of *MHC-restriction* is essential in narrowing the massive genetic diversity of the unselected TCR repertoire. Remarkably, while there is combinatorial diversity (which Vα and Vβ gene segments are incorporated into a given TCR), the truly specialized nature of a given TCR must stem from the RAG1/RAG2 mechanism of somatic DNA rearrangements which focus structural diversity to the CDR3α and CDR3β loop amino acid (a.a.) sequences [6, 12, 19]. Thus, it follows that the capacity of this genetic system for clonotypic structures in terms of CDR3α/CDR3β loops must entail the fine selection of a population of TCR-bearing clones that are best suited to protect the host. The same teleological considerations apply to the sIg on B cells, but their ligands are quite different. Whereas sIg can vary over five orders of magnitude in binding equilibrium constants with both various soluble proteins and insoluble proteins (e.g., bacterial toxins, or proteins on the surface of a bacterium or virus, respectively), TCR have been shown to share micromolar binding constants, despite vast differences in the a.a. sequences of their CDR3 loops, and indeed in the different pMHC ligands they recognize [17, 20, 21]. Clearly however, some pathogens favor MHC-I dominated responses, e.g., direct infected-cell killing by CD8 T cells is a hallmark of the response to viral pathogens, while extra-cellular pathogens (e.g., most *E. coli*) favor MHC-II effector functions, e.g., production of cytotoxic immunoglobulins (ref. 1).

One clue as to how the TCR functions might be in how *mechanical* forces have apparently become intertwined in this system of trans-genetic, protein structural recognition [13–17]. More specifically, we have become interested in the role of TCR *dynamics* at the initial stage of the recognition process [11]. Here, we build upon previous derivation of a rotational dynamics, multivariable calculus approach used to model TCR V-domain binding to the conspicuous, highly conserved, inclined-plane (IP) structures of the two opposing α-helices prominent in MHC-I and MHC-II proteins [22, 23]. A dichotomy was revealed by the calculations of these dynamics for Vα versus Vβ binding pMHC-I versus pMHC-II ligands. As such, a novel path is put forward for understanding evolved T-cell functions; this theory entails prevailing evidence that relative TCR binding affinity is not the mechanism of TCR antigen specificity [11, 16, 17, 20, 24].

## Results

Previous work defined a clonotypic *volume element* (*dV*) for each V-domain in TCR:pMHC complexes utilizing standard conversion to spherical coordinates and a general triple-integral equation. Here, we used the same analytical geometry and calculus approach to estimate the *torque* imparted by the apparent 45° rotation of each V-domain. Why 45°? Because the comparative geometry showed that the CDR2-loop closest contact with its cognate MHC α-helix defines a range (between different TCR:pMHC structures) corresponding to an arc distance of the central cysteine rotating through exactly 45° [11]. As originally hypothesized [25], the rotations for both domains are in the counter-clockwise (cc) direction, because the two, conspicuous IP of the MHC α-helices face in *opposite* directions (e.g., easily seen in Fig. 2C). Thus, as appreciated by the so-called *right-hand rule*, we can see that when both V-domains rotate *cc* that the direction of the torque force for both domains would be toward the T cell (ref. 26). This is intuitive if, the effect of this torque is to move each V-domain up to the top of their cognate MHC α-helix IP (Fig. 1-Fig. 4).

**Figure 1.**
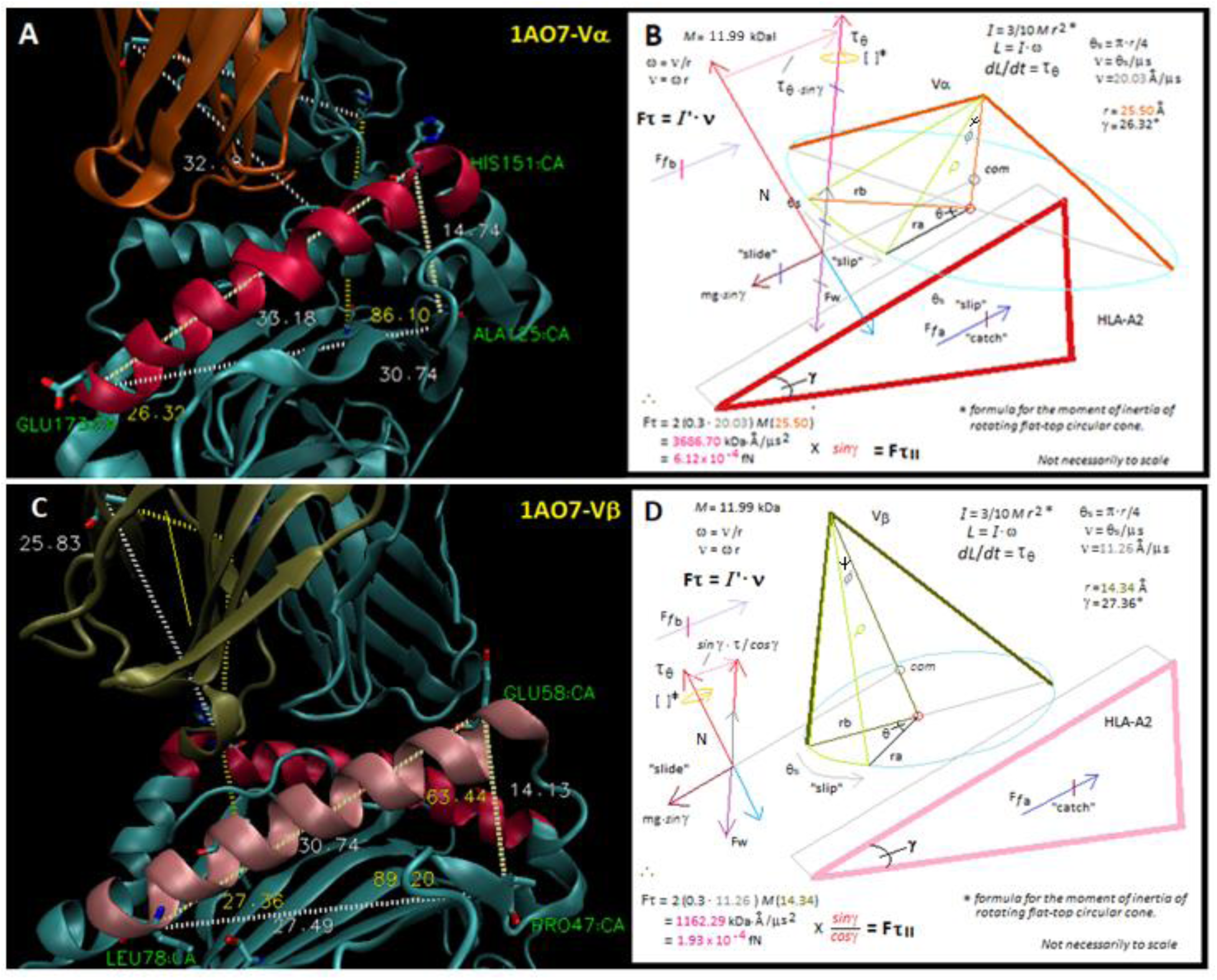
Vector analysis of human TCR:pMHC class-I structure 1AO7 highlighting Vα (A & B) and Vβ (C & D) interactions with the inclined planes of the A2 α-2 helix and A2 α-1 helix, respectively. (A) TCR Vα shown in *burnt orange*, HLA-A2 α-2 helix shown in *red*. A large right triangle is defined by the near 90° angle H151:A125:E173. The γ-angle is defined by A125:E173:H151 (= 26.32°). (C) TCR Vβ shown in *tan*, HLA-A2 α-1 helix shown in *pink*. A large right triangle is defined by the near 90° angle E58:P47:L78. The γ-angle is defined by P47:L78:E58 (= 27.36°). The Vα and Vβ cones defined by spherical coordinates in the triple integral equation (ref. 11) are shown as the downward projection (B & D), where rotation through 45° specifies the *dV* volume elements (*lime* cone slices). The calculation for *torque* is in (B & D), where standard vector analysis for an object on an inclined plane is shown [26, 27]. Note that the torque force parallel to the IP is calculated by trigonometry and is countered by the sliding force. H-bonds including those along the α-helix with the CDR2 loop create a modeled frictional force involving the V-domain at a characteristic distance from the apex of the IP. 1AO7 *PDB* file available at *NCBI* (http://www.ncbi.nlm.nih.gov).

**Figure 2.**
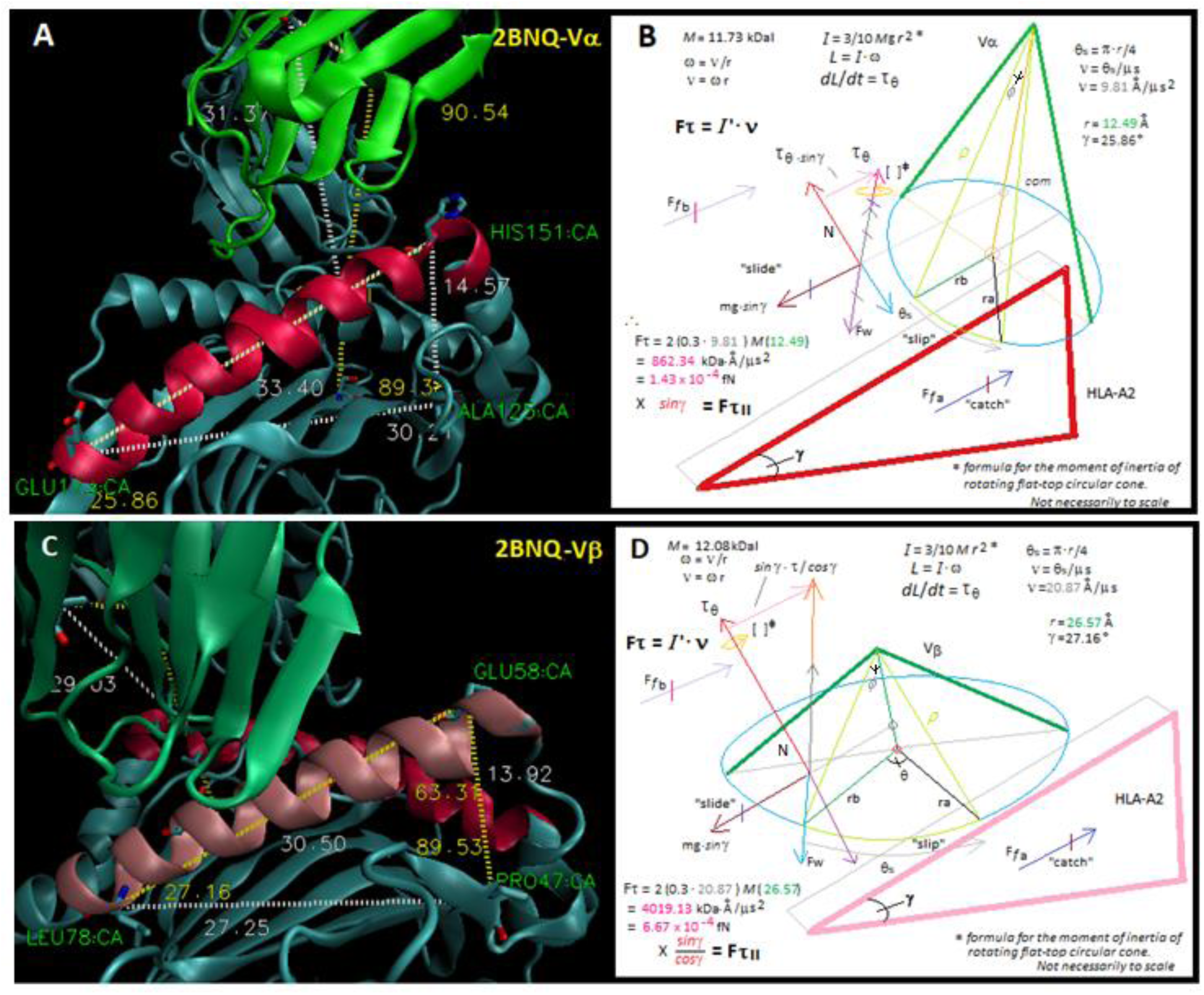
Vector analysis of human TCR:pMHC class-I structure 2BNQ highlighting Vα (A & B) and Vβ (C & D) interactions with the inclined planes of the A2 α-2 helix and A2 α-1 helix, respectively. (A) TCR Vα shown in *green*, HLA-A2 α-2 helix shown in *red*. A large right triangle is defined by the near 90° angle H151:A125:E173. The γ-angle is defined by A125:E173:H151 (= 25.86°). (C) TCR Vβ shown in *green3*, HLA-A2 α-1 helix shown in *pink*. A large right triangle is defined by the near 90° angle E58:P47:L78. The γ-angle is defined by P47:L78:E58 (= 27.16°). The Vα and Vβ cones defined by spherical coordinates in the triple integral equation (ref. 11) are shown as the downward projection (B & D), where rotation through 45° specifies the *dV* volume elements (*lime* cone slices). The calculation for *torque* is in (B & D), where standard vector analysis for an object on an inclined plane is shown [26, 27]. Note that the torque force parallel to the IP is calculated by trigonometry and is countered by the sliding force. H-bonds including those along the α-helix with the CDR2 loop create a modeled frictional force involving the V-domain at a characteristic distance from the apex of the IP. 2BNQ *PDB* file available at *NCBI* (http://www.ncbi.nlm.nih.gov).

**Figure 3.**
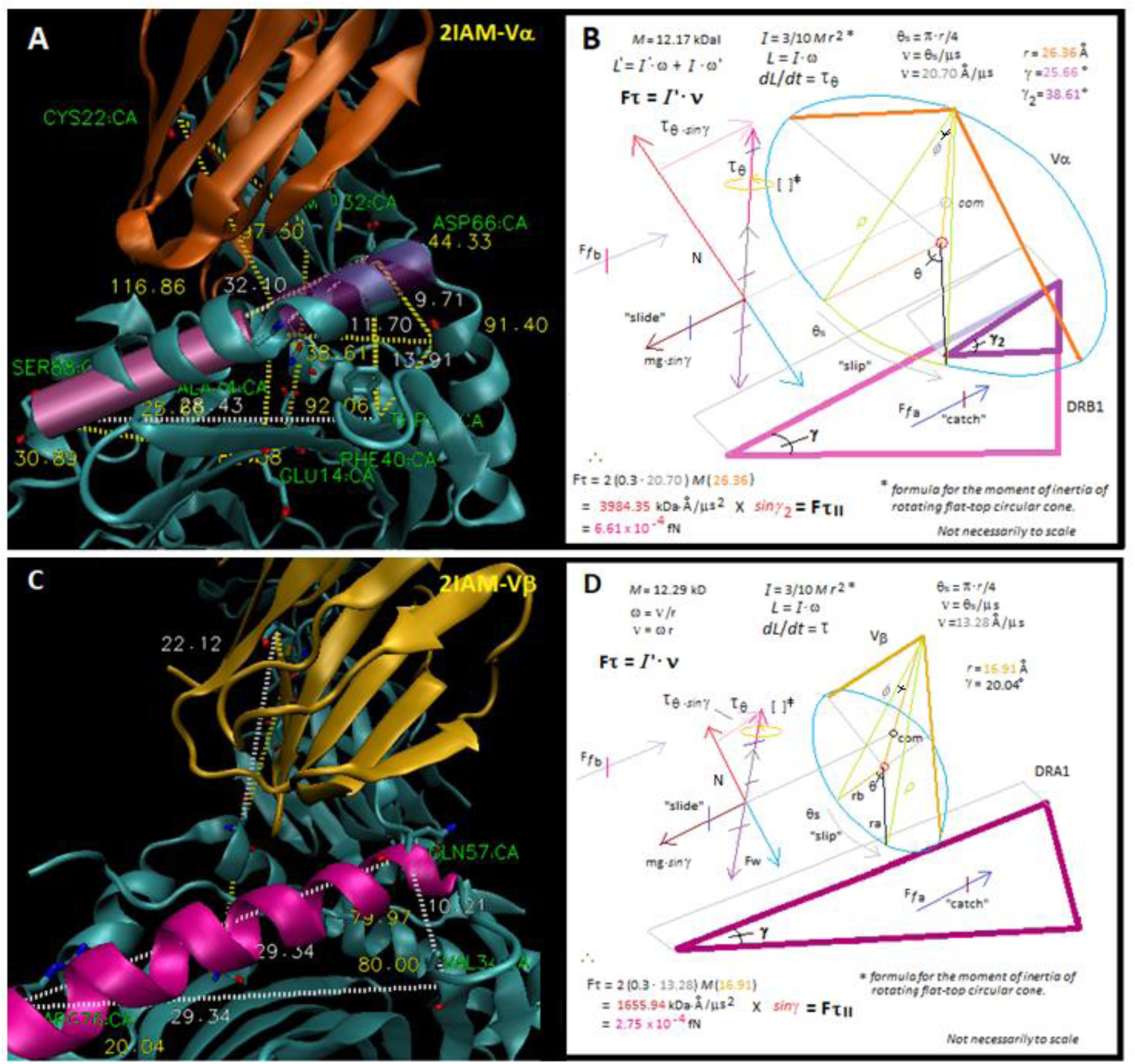
Vector analysis of human TCR:pMHC class-II structure 2IAM highlighting Vα (A & B) and Vβ (C & D) interactions with the inclined planes of the DRB1 and DRA α-helices, respectively. (A) TCR Vα shown in *orange*, HLA-DRB1 α-helix shown in *purple* and *mauve*. A large right triangle is defined by the near 90° angle D66:F40:S88. The γ_2_-angle of the small right triangle (*purple*) derives parallel torque (= 38.61°). (C) TCR Vβ shown in *gold*, HLA-DRA α-helix shown in *magenta*. A large right triangle is defined by the near 90° angle Q57:V34:R76. The γ-angle is defined by V34:R76:Q57 (= 20.04°). The Vα and Vβ cones defined by spherical coordinates in the triple integral equation (ref. 11) are shown as the downward projection (B & D), where rotation through 45° specifies the *dV* volume elements (*lime* cone slices). The calculation for *torque* is in (B & D), where standard vector analysis for an object on an inclined plane is shown [26, 27]. Note that the torque force parallel to the IP is calculated by trigonometry and is countered by the sliding force. H-bonds including those along the α-helix with the CDR2 loop create a modeled frictional force involving the V-domain at a characteristic distance from the apex of the IP. 2IAM *PDB* file available at *NCBI* (http://www.ncbi.nlm.nih.gov).

**Figure 4.**
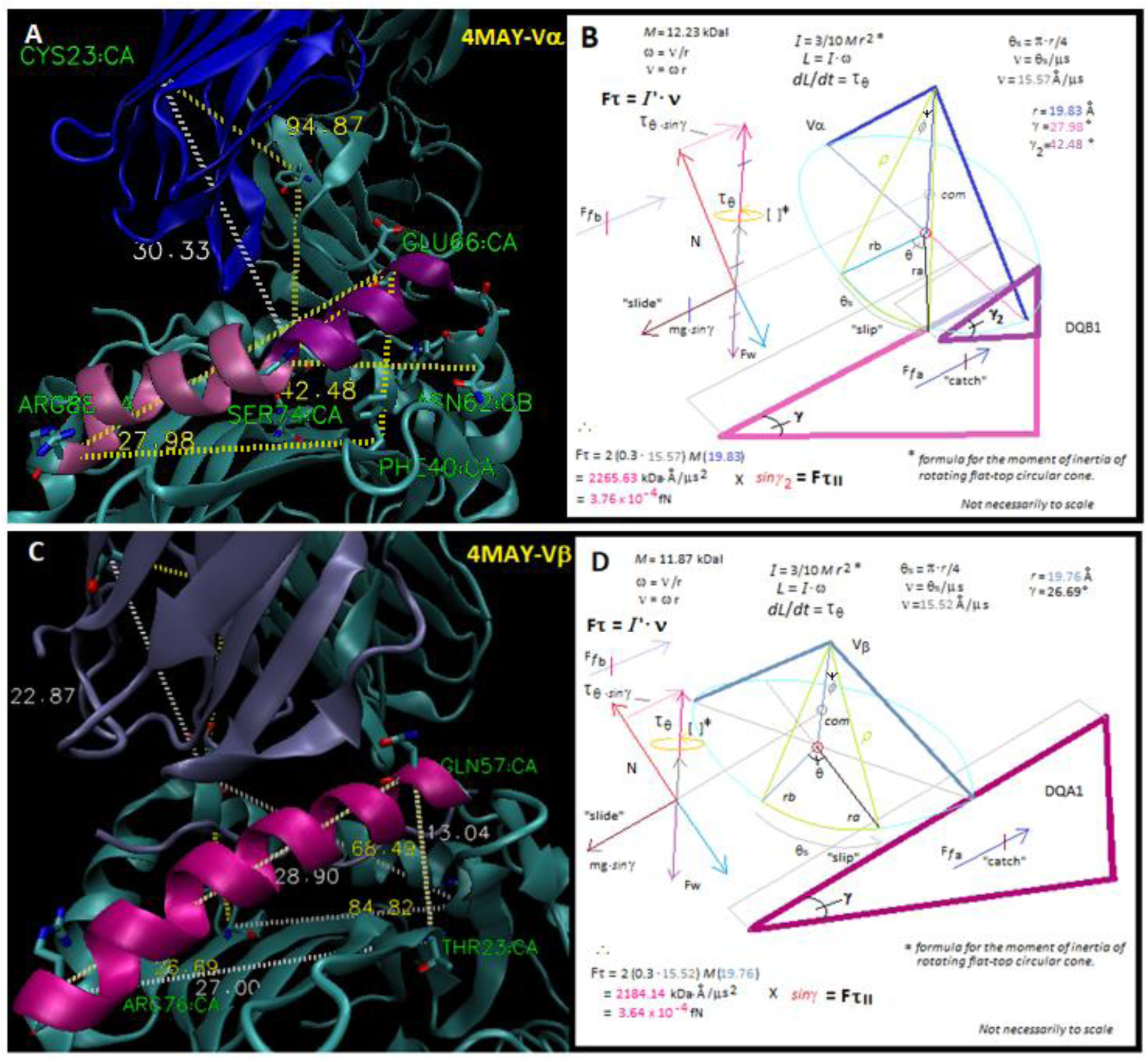
Vector analysis of human TCR:pMHC class-II structure 4MAY highlighting Vα (A & B) and Vβ (C & D) interactions with the inclined planes of the DQB1 and DQA1 α-helices, respectively. (A) TCR Vα shown in *dark blue*, HLA-DQB1 α-helix shown in *purple* and *mauve*. A large right triangle is defined by the near 90° angle E66:F40:R88. The γ_2_-angle of the small right triangle (*purple*) derives parallel torque (= 37.70°). (C) TCR Vβ shown in *ice blue*, HLA-DQA1 α-helix shown in *magenta*. A large right triangle is defined by the near 90° angle Q57:T23:R76. The γ-angle is defined by T23:R76:Q57 (= 26.69°). The Vα and Vβ cones defined by spherical coordinates in the triple integral equation (ref. 11) are shown as the downward projection (B & D), where rotation through 45° specifies the *dV* volume elements (*lime* cone slices). The calculation for *torque* is in (B & D), where standard vector analysis for an object on an inclined plane is shown [26, 27]. Note that the torque force parallel to the IP is calculated by trigonometry and is countered by the sliding force. H-bonds including those along the α-helix with the CDR2 loop create a modeled frictional force involving the V-domain at a characteristic distance from the apex of the IP. 4MAY *PDB* file available at *NCBI* (http://www.ncbi.nlm.nih.gov).

### Estimation of V-domain torque

The equation series used to calculate a particular V-domain *torque* (*τ*_θ_) is shown in Figure 1B & D through Figure 4B & D, where the general equations are described below. Firstly, the basic parameters for each were measured from the appropriate *PDB* file in *VMD* software (https://www.ks.uiuc.edu/) using fixed a.a. positions for each variable, as previously described [11, 25]. Briefly, mass (*M*) was computed from the a.a. sequence of each V-domain starting at position-1 in the crystal structure through the F-G-R-G motif near the end of the ≈ 110 a.a. domain using the *ExPASy ProtParam* algorithm (https://web.expasy.org/). The distance *rho* (*ρ*) can be seen to partially specify the shape of a ‘cone-slice’ volume element (*dV*) in Figure 1 through Figure 4 (*lime green* cone slice), where the triple-integral equation [11] is shown in the captions to Table 1 and Table 2 (see also, **Materials and Methods**). Crucially, the distance, rho (in angstroms, Å), is used in the trigonometric determination of the *radius* (*r*) of each uniquely shaped V-domain cone: ***r*** = *sin ϕ* (*ρ*) (ref. 27).

**Table 1.**
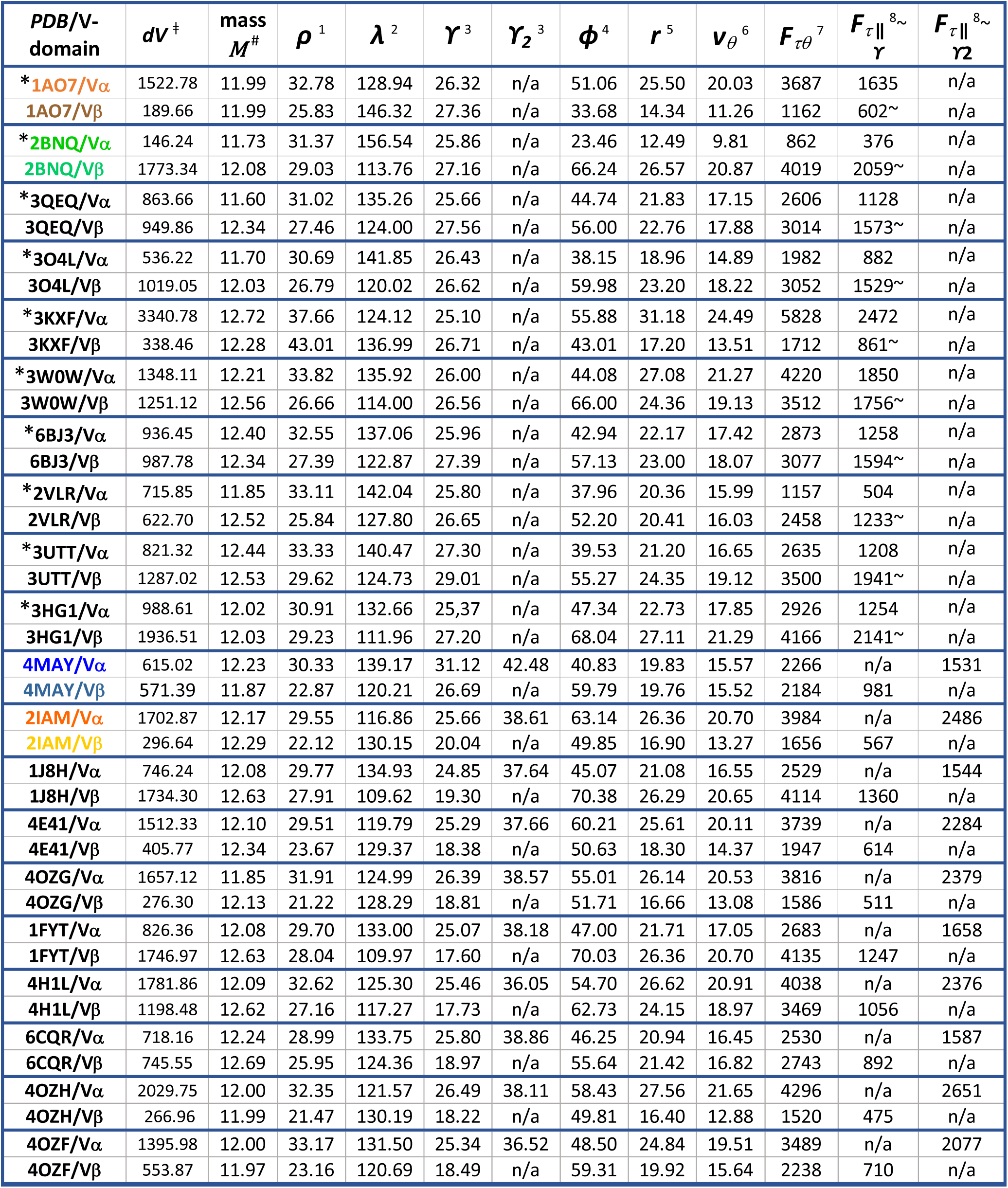
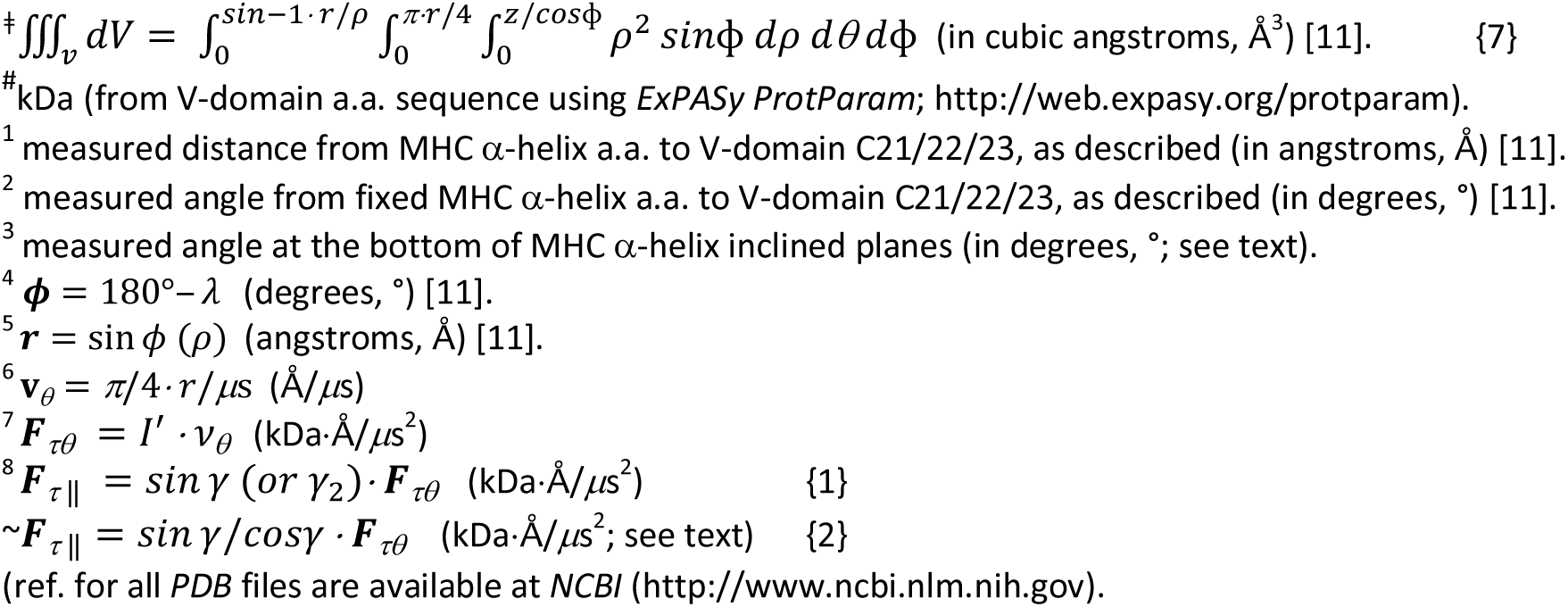
Summary of calculated V-domain rotational dynamics of TCR:pMHC-I * & TCR:pMHC-II solved structures.

**Table 2.**
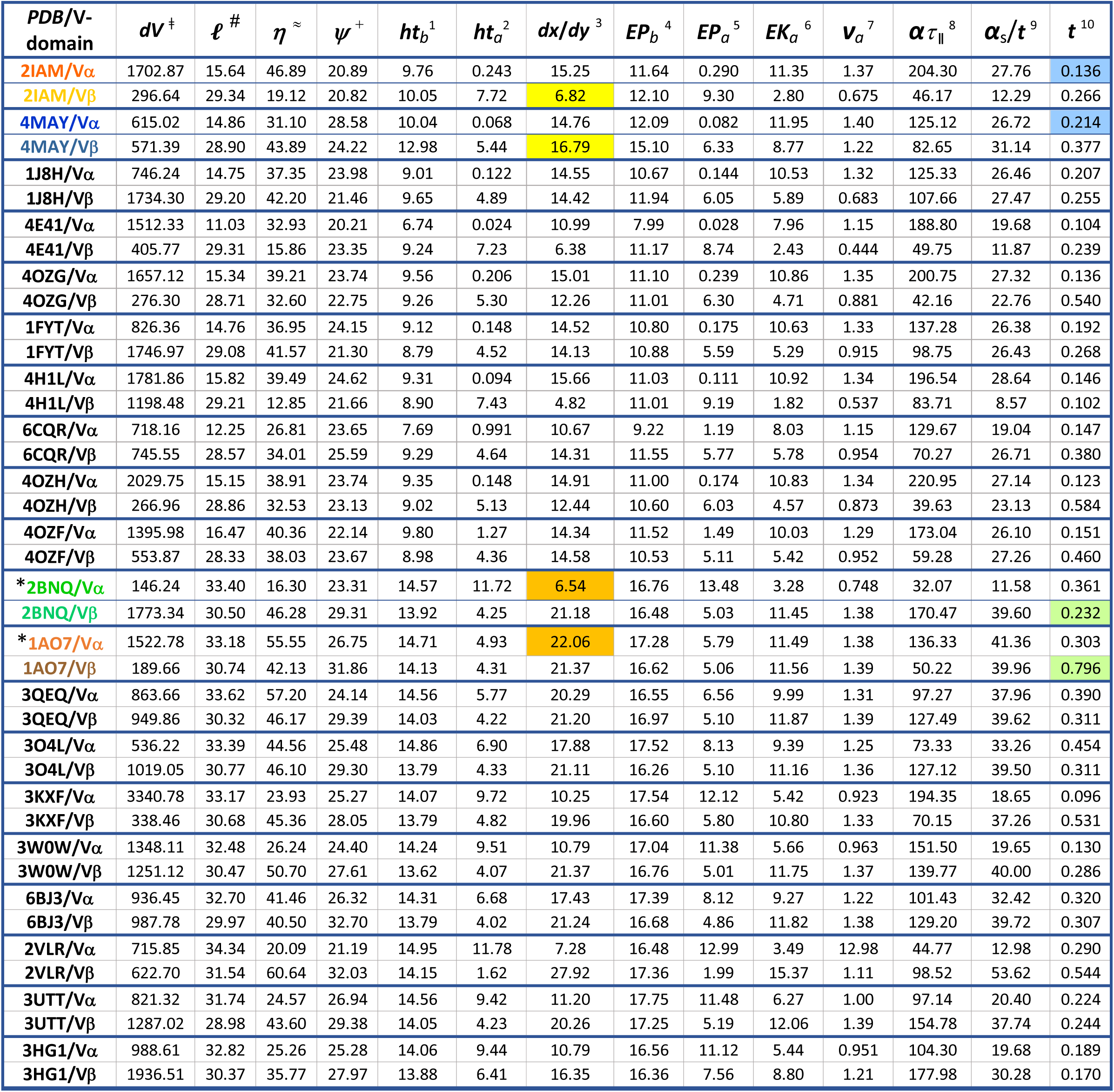
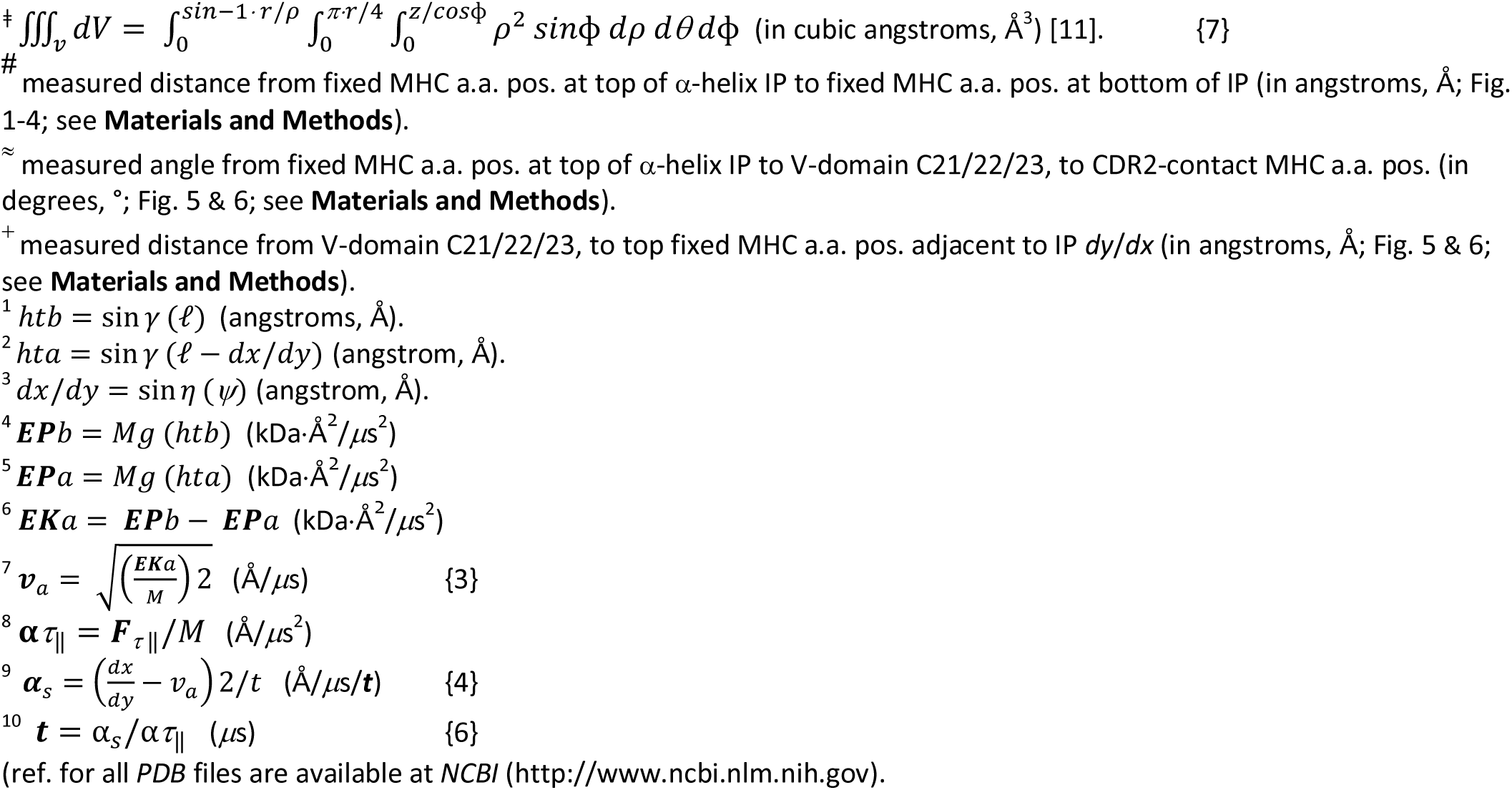
V-domain *slide-time* on the MHC α-helix inclined planes of TCR:pMHC-II & TCR:pMHC-I* complexes.

To estimate the *arc* distance path of CDR2, rotation of (*r*) by 45° gives the *dV* arc distance in angstroms, Å (*θ*_s_) by the equation: *θ_s_* = *π*/4 · *r*; see ‘*r* -*before*’ (***rb***) rotation to ‘*r* -*after*’ (***ra***) in Figure 1B & D-4B & D. This leads to an assumption on *velocity* for the rotation, in that we assume this takes 1.00 *μ*s, i.e., ***v****_θ_* = *π*/4 · *r*/*μ*s (in Å /*μ*s). The general equation: ***I*** = (3/10) *M*·*r*^2^ for the moment of inertia (*I*) of a rotating flat-top cone, based upon the common relative position for center-of-mass (*com*), was used to find the first derivative: ***dL*/*dt =*** *τ_θ_* for the angular momentum *rate-of-change*. This equation reduces to: ***F*** *τ_θ_* = *I* ′ ·*v_θ_*, which defines the *torque* force in the direction shown by the *magenta* vectors in each figure [26, 27].

Note, that to find the component of torque parallel (‖) to each IP, it was clear that the force vector is usually opposite ***F***w (e.g., as in Fig. 1A & B) and thus the equation:

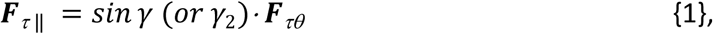

could be used, i.e., where the angle, (γ) or (γ_2_) is the angle at the bottom of the particular MHC α-helix IP ‒ always using fixed a.a. positions between similar MHC class and helix (Fig. 1-Fig. 4; seen also in, Fig. 5 & 6). Here, the (γ_2_) angle is critical. This angle is made by the split of the long α-helix into two separated α-helices, i.e., creating a top IP at the apex of MHC-II beta chain α-helices (Fig. 3A & B, Fig. 4A & B, Fig. 6A & 6B). Notably, the direction of the overall torque is sometimes perpendicular to the slope of the IP (Fig. 1C & D, Fig. 2C & D; Table 1 & 2); only seen for the Vβ domain interaction with the MHC-I α-1 helix. In this case, the component of the torque force parallel to the slope of the IP can be calculated by the equation:

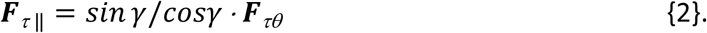

**Figure 5.**
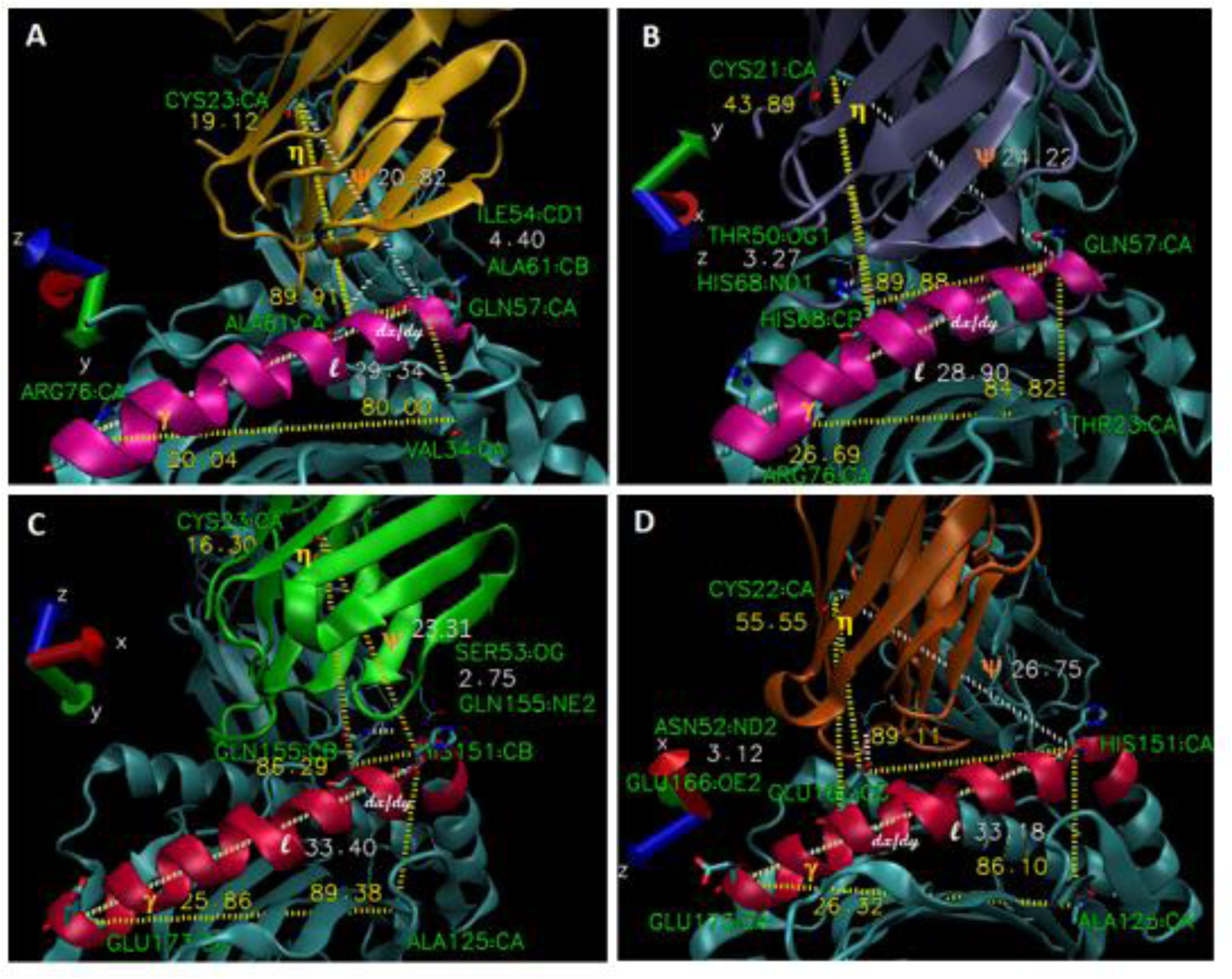
Variability in the inclined plane position of V-domains in TCR:pMHC-II and TCR:pMHC-I complexes. (A) 2IAM-Vβ domain (highlighted in *gold*) interaction with DRA α-helix (*magenta*). (B) 4MAY-Vβ domain (*ice blue*) interaction with DQA1 α-helix (*magenta*). (C) 2BNQ-Vα domain (*green*) highlighted interaction with HLA-A2 α-2 helix (*red*). (D) 1AO7-Vα domain (*burnt orange*) interaction with HLA-A2 α-2 helix (*red*). Key a.a. defining the near 90° right triangle of the inclined planes shown in licorice by atom (names in *green*); angles shown in *yellow*, measured in *VMD* (yellow-dotted lines). The γ-angle used in the equation series defining ‘slide time’ is shown in *yellow*, near the *melon* coloured ‘γ’ symbols. The near 90° angle from the IP apex a.a. to the α-helix a.a. contacting the CDR2 loop to the central cysteine is shown, where the angle from this a.a. to the central cysteine to the apex a.a. defines the *η*-angle (*yellow* ‘η’ symbol) used to calculate the distance *dx*/*dy* (*white* annotation) by trigonometry. The full IP distance (l) was measured by *VMD bond-label* tool (shown in *white* and white-dotted lines). The CDR2-loop contact with the α-helix is shown by the short, white-dotted bond lines and the value is in *white* between the a.a. atoms in the contact (shown in *green*). The distance from the central cysteine to the IP apex a.a. (*ψ*) is shown by a white-dotted line near the ‘ψ” symbol (*orange*). The remainder of structures are in *cyan*. View orientation for each *PDB* is shown by the *xyz-axis* tool. *PDB* files available at *NCBI* (http://www.ncbi.nlm.nih.gov).

**Figure 6.**
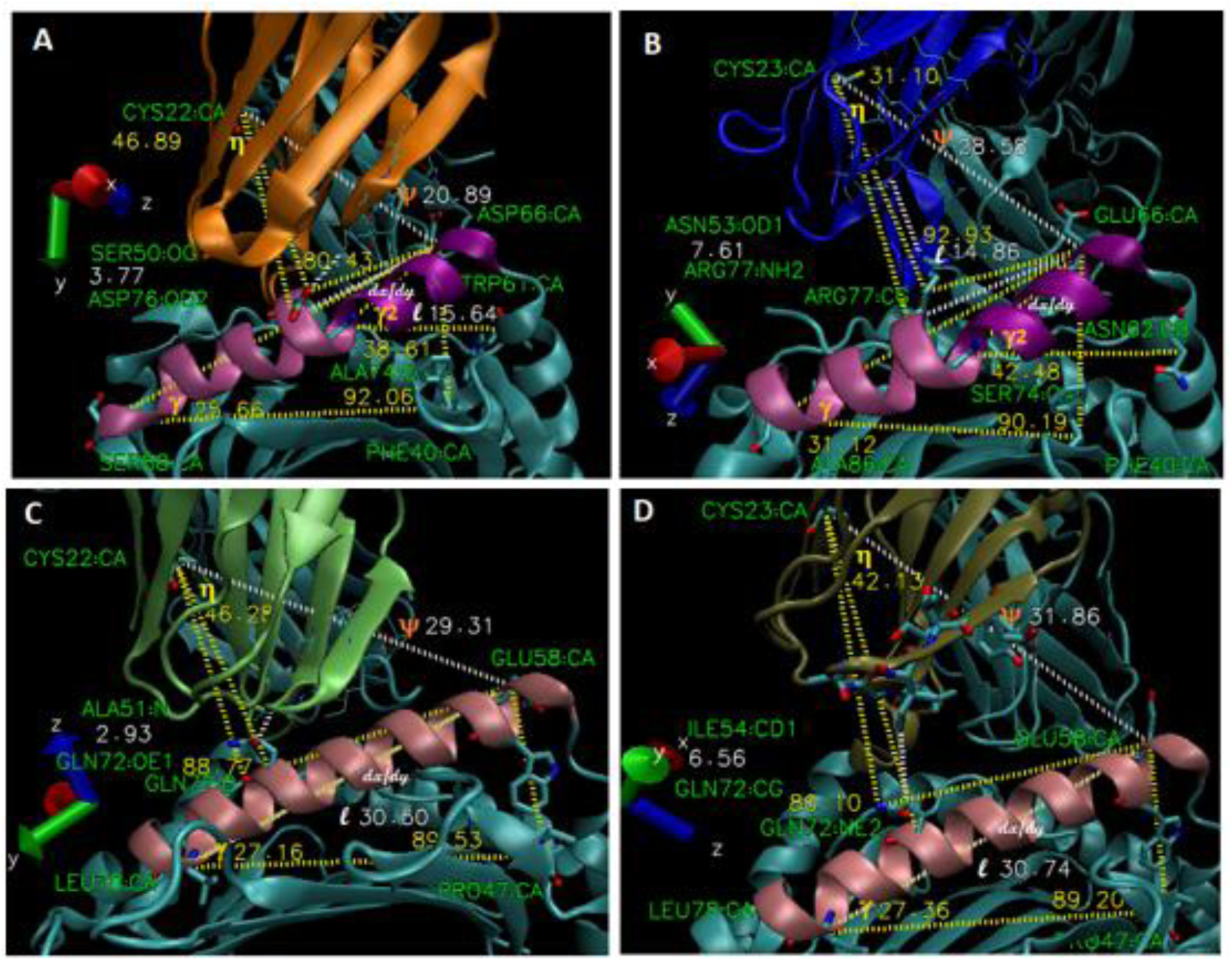
Consistency in the inclined plane position of V-domains in TCR:pMHC-II and TCR:pMHC-I complexes. (A) 2IAM-Vα domain (highlighted in *orange*) interaction with DRB1 α-helix (*purple and mauve*). (B) 4MAY-Vα domain (*dark blue*) interaction with DQB1 α-helix (*purple and mauve*). (C) 2BNQ-Vβ domain (*lime*) highlighted interaction with HLA-A2 α-1 helix (*pink*). (D) 1AO7-Vβ domain (*tan*) interaction with HLA-A2 α-1 helix (*pink*). Key a.a. defining the near 90° right triangles of the inclined planes shown in licorice by atom (names in *green*); angles shown in *yellow*, measured in *VMD* (yellow-dotted lines). The γ-angle used in the equation series defining ‘slide time’ is shown in *yellow*, near the *melon* coloured ‘γ’ symbols. Note that MHC-II beta chains have a split IP, where a smaller right triangles’ γ_2_-angle is measured using an adjacent N-term. a.a. to draw a vector that bisects (perpendicular) the larger triangles’ height vector (yellow-dotted lines). This γ_2_-angle is shown in *yellow* near the *melon* coloured ‘γ_2_’ symbols, which were used in the equation series for Vα:MHC-II beta interactions. The near 90° angle from the IP apex a.a. to the α-helix a.a. contacting the CDR2 loop to the central cysteine is shown, where the angle from this a.a. to the central cysteine to the apex a.a. defines the *η*-angle (*yellow* ‘η’ symbol) used to calculate the distance *dx*/*dy* (*white* annotation) by trigonometry. The full IP distance (l) was measured by *VMD bond-label* tool (shown in *white* and white-dotted lines). Note this distance is for the smaller right triangle for MHC-II beta α-helices. The CDR2-loop contact with the α-helix is shown by the short, white-dotted bond lines and the value is in *white* between the a.a. atoms in the contact (shown in *green*). The distance from the central cysteine to the IP apex a.a. (*ψ*) is shown by a white-dotted line near the ‘ψ’ symbol (*orange*). The remainder of structures are in *cyan*. View orientation for each *PDB* is shown by the *xyz-axis* tool. *PDB* files available at *NCBI* (http://www.ncbi.nlm.nih.gov).

As shown in Table 1, the IP-parallel component of torque showed a broad range from 376 kDa·Å/*μ*s^2^ (2BNQ Vα) to 2651 kDa·Å/*μ*s^2^ (4OZH Vα). It is clear that this is not due to differences in V-domain *mass* (all are similar, Table 1), but rather it is the volume element (*dV*), partially defined by the radius of the cone shape, which is clonotypic. The radius value also determines the calculated velocity through the volume swept by the *dθ* derivative; more fundamentally, (*r*) is based on only two variables that both describe the relative location of the V-domain central cysteine (C21/22/23) in space relative to the pMHC groove [11].

By trigonometry: ***r*** = sin *ϕ* (*ρ*), thus (*r*) depends on the angle *phi* and the distance rho. As previously described [11, 25], the phi angle is the difference between the theoretical normal (180°) and the ‘tilt’ angle, which was measured in *VMD* for all these structures. Specifically, *tilt* (λ) is the angle formed by a line segment running from a fixed (β-sheet) a.a. at the bottom of the groove to a fixed a.a. of the α-helix directly above it, then another line segment from this α-helix a.a. to the central cysteine (e.g., F40(DQB1):E66(DQB1):C21(Vβ), for 4MAY Vβ = 120.21°; Table 1). Thus, the tilt angle determines phi *ϕ =* 180 – *λ*. The distance rho is the measured distance in each structure between the fixed a.a. of the α-helix (for 4MAY Vβ it is E66) and the central cysteine. This indicates that one can use the location in space of any V-domain’s central cysteine relative to its pMHC-groove to describe a mathematical model yielding a clonotypic cone representing perhaps key geometry in the structure.

Moreover, mathematically modeled *rotation* of this cone through 45° in each structure yields strikingly different volume elements through a path on its cognate MHC α-helix, which as stated, correlates with the range of closest CDR2 contacts between different structures (≈ 10 a.a. range) [11]. Here, we show that these variables can be further used to estimate the *torque* of any given V-domain (see below, Table 1, and **Discussion**). In summary (Table 1), so-called ‘highly-restricted’ *dV*, for which the volume element (Å^3^) is reduced by ≥ 70% (*n* = 40, mean: 1058.94, SD: 642.39) could be found for both Vα (2BNQ) and Vβ (1AO7, 2IAM, 4OZH, 4OZG), occur across MHC classes, and there was the expected linear relationship between *dV* and torque (**α***τ*_∥_); see **Supplement I** for the plots, where ***R***^2^ = 0.985, 0.864, 0.805, 0.954 for Vα (cl II), Vβ (cl II), Vα (cl I), and Vβ (cl I), respectively (*n* = 10).

### Estimation of V-domain *slide time* on MHC α-helix IP

Since V-domain torque through the CDR2-MHC α-helix binding path (*dθ*) correlates in *direction* with “lifting” the V-domain from an initial position to the apex of the MHC α-helix IP, the next task was to use *energy conservation* with respect to an object on an inclined plane [26, 27] to investigate *magnitude* of the torque vector, and to solve for the *time* indicated by these mechanics, i.e., how long it would take each V-domain to ‘slide’ to the position in the ground-state structure (Fig. 1B & D-Fig. 4B & D). Recall, that we assume a single *μ*s for velocity. How long it would take a V-domain to actually rotate through the 45° indicated in the model is certainly not known (see **Discussion**). However, with this assumption, the goal was to normalize the velocity variable, such that differences in torque, and indeed, ‘slide’ could be isolated. As shown in Figure 5 and 6, a fixed a.a. position at the top of each MHC α-helix type was used to measure the best right angle possible with the CDR2 loop’s closest contact MHC position along the α-helix (shown as a short, white-dotted bond line). For example, in 4MAY the bond length is 3.27 Å for Vβ, where T50:OG1 hydrogen-bonds with the HLA-DQA1 chain α-helix via H68:ND1 (Fig. 5B). Any atom of this MHC position was chosen to obtain the best 90° angle by the *VMD angle-label* tool; this is to show how close to a right angle these relationships are in the ground-state structures (e.g., for 4MAY-Vβ, 89.88°). To estimate the angle (*η*), which runs from the closest CDR2-contact α-helix a.a. (Cα) to the central cysteine (Cα), then to the IP apex a.a., Cα (e.g., Q57 for DRA and DQA1 helices, Fig. 5A & 5B; H151 for A2 α-2 helix, 5C & 5D) was measured with *VMD* (yellow dotted line to white dotted line angle near the *yellow* ‘η’ markers). The distance from the apex a.a. to the central cysteine was measured and is shown next to the white-dotted bond line, i.e., see the *orange* ‘ψ’ symbols. Thus, the trigonometry is: ***dx***/***dy*** *sin η* (*ψ*), and corresponds to the distance in Å traveled by the CDR2-loop contact with the MHC α-helix (Fig. 5 & 6). Next, in order to estimate the *potential energy* of an object on an IP, it is necessary to know the *height* of a given position along the slope [26]. For the ‘before’ position, the apex a.a. of the α-helix to a fixed β-sheet a.a. in each type of MHC (e.g., 4MAY-Vβ is T23; Fig. 5B) to a similarly fixed a.a. at the lowest IP position (e.g., 4MAY-Vβ is R76; Fig. 5B) defines a near 90° angle in all the structures (Fig. 5 & 6). Thus, ‘height before’ (*htb*) is calculated by: ***htb*** = *sin γ* (*ℓ*) from the angle (γ) and the measured bond length between the apex a.a. Cα and the lowest a.a. position of the IP Cα; this is the length (*ℓ*), shown in *white* next to the white-dotted bond line. In Figure 6, is shown how the upper separate IP for the MHC-II beta-chain α-helix, *htb*, was determined by using an adjacent a.a. position, N-term. to the apex a.a. to estimate this bottom of the IP angle, i.e., with a vector from the adjacent a.a. (N62:CB for 4MAY; Fig. 6B) that bisects perpendicular to the height vector of the larger right triangle defining the full-length IP. To obtain the height at the point along the IP corresponding to the CDR2 contact with MHC/position of the central cysteine, i.e., the ‘height after’ (*hta*), we used the CDR2-contact position in the ground-state. Thus, the distance of the full IP minus *dx/dy* is the hypotenuse of a right triangle, where the height-side of the triangle is *hta*, by: ***hta*** = *sin γ* (*ℓ* − *dx*/*dy*). From these heights we can calculate *potential energy* at the two positions on the IP: ***EP****b = Mg* (*htb*), and ***EP****a = Mg* (*hta*), respectively. Further, *kinetic energy* at *hta* can then be determined from the *first law*: ***EK****a = **EP**b* − ***EP****a*. Next, from the kinetic energy we can calculate *velocity* at *hta*, solving for (***ν*** *_a_*) in the equation:

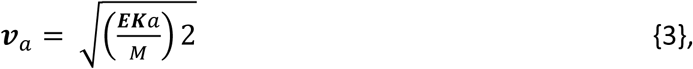

then, we use the third kinematic formula for acceleration (**α**):

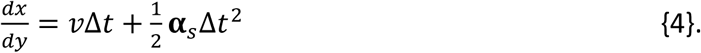

Where, we solve for **α** _s_, which is the value for V-domain *acceleration* down the IP. Importantly, the answer is expressed as Å/*μ*s/*t*, such that the variable ‘***t****’* can be isolated, when (for a stationary object on an IP) the acceleration parallel-up the plane must equal the acceleration parallel-down the plane [26, 27]. Specifically:

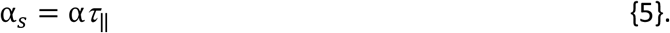

The torque (***F****_τ_*_∥_) was determined as previously indicated (Fig. 1-Fig. 4), and *Newton’s second law*: ***F*** = *M*ατ_∥_ was used to find the acceleration ‘up’ the IP. Again, the ‘slide-time’ down the IP must allow for equal acceleration up and down the IP, and could be calculated by:

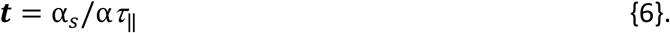

Two solutions to this series of equations for two sample V-domains are shown below; solutions for highlighted V-domain values reported in Table 2 are available in **Supplement I**. A broader sample of MHC-I and MHC-II restricted TCR further showing this MHC class relationship on positioning of CDR2-loop contacts along cognate α-helices is available (*n* = 40) in Table 3A & 3B, **Supplement I**.

### Sample ‘slide-time’ solutions*

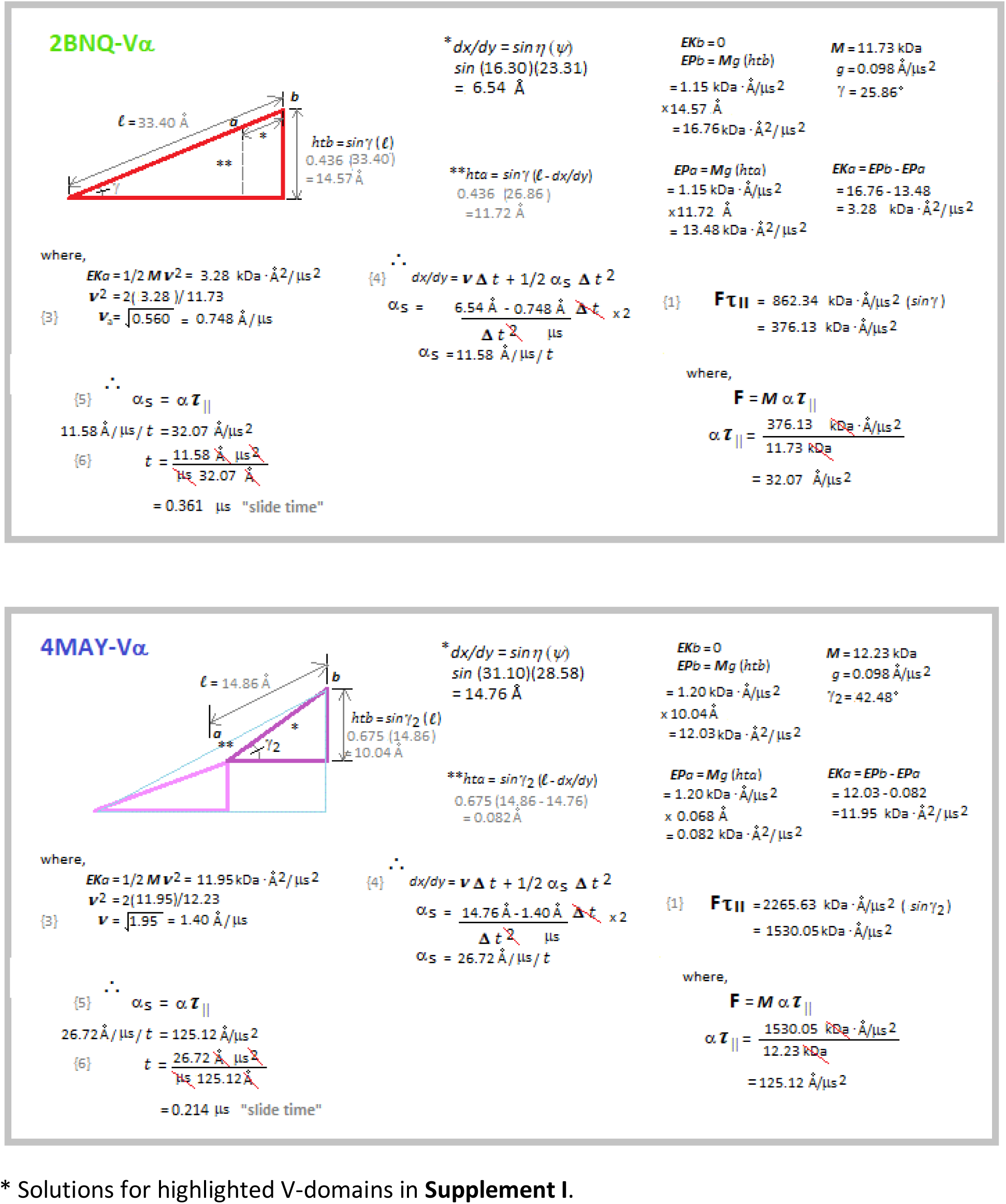

### Dichotomy of V-domain positioning on MHC-II versus MHC-I inclined planes

As indicated, the α-helix of human MHC-II beta chains is split into two separate inclined planes (Fig. 3A & B, Fig. 4A & B, Fig. 6A & 6B). The large right triangle specifying a full IP, is easily observed and measured in *VMD* for the MHC-II alpha chain α-helix (Q57(apex IP):T23(DQB1) or V34(DRB1), β-sheet floor:R76(bottom a.a. of the IP); see Fig. 3C & D, Fig. 4C & D, Fig. 5A & 5B); also note, DPA5 is similar (*PDB* 3WEX; ref 28), where the ≈ 90° angle runs Q57:M23:R76 at 83.10°. However, as in HLA-DQB1 and HLA-DRB1, there is a split in this IP for DPB5 as well. For DPB5 the large triangle runs D64:F38:A86 at 94.14°. The smaller apex right triangle is estimated (as for DR and DQ, see previous) by using an adjacent N-term. a.a. position giving the IP angle for the smaller triangle, which is steeper at ≈ 39° (MHC-II beta) than the larger triangle. For DPB5 this angle is N60:V72:D64 at 36.98° (data not shown). Importantly, both IP of MHC-I are the full-length type (i.e., like MHC-II α chains), e.g., see Figure 1 and Figure 2. There is consistent placement of the TCR CDR2 contact with the MHC-II beta chain α-helix at the junction between the top helix IP and the bottom IP; that is, between different TCR:pMHC-II structures (Table 2 & **Supplement I**). Yet, that is not the case for ‘time’ of the Vα domain interaction as calculated with the MHC-II beta α-helix, see below (Fig. 6A & 6B; Table 3B & **Supplement I**). The positioning of the Vα CDR2 contact (*dx/dy*) is 15.25 Å for the DR-restricted 2IAM-Vα and 14.76 Å for the DQ-restricted 4MAY-Vα (Table 2). By contrast, Vβ interaction with the MHC-II alpha IP is variable, e.g., *dx/dy* for 4MAY-Vβ is 16.79 Å, 2IAM-Vβ is 6.82 Å (Table 2, highlighted in *yellow*; see also Table 3B & **Supplement I**). Conspicuously, for MHC-I restricted TCR the opposite is seen, i.e., Vα interaction with the heavy-chain α-2 helix is variable in terms of positioning of the CDR2 contact along the helix (1AO7-Vα *dx*/*dy* is 22.06 Å, where 2BNQ-Vα is 6.54 Å; Table 2, highlighted in *orange*; see also Table 3A & **Supplement I**). Meanwhile, Vβ is the consistent interaction with the α-1 helix, e.g., 1AO7-Vβ *dx*/*dy* = 21.37 Å and 2BNQ-Vβ *dx*/*dy* = 21.18 Å. Importantly, embedded in this dichotomy is the finding that ‘variability’ for the V-domains that show consistent mobility on the IP (Vα/MHC-II beta; Vβ/MHC-I α-1) was observed in the *time* spent sliding on the IP (Table 2, highlighted in *blue* and *green*). By inspection, we find the dichotomy to be consistently observed between the TCR:pMHC-II and TCR:pMHC-I structures (Table 2; Table 3A & 3B, & **Supplement I**). Some Vβ on MHC-I α-1 helix show ‘slippage’ past Q72, e.g., *PDB* 2VLR (see above). With relatively few exceptions (*n* = 40), it is interesting to find this dichotomy of the two V-domains positioning on the two α-helices of the two MHC classes, i.e., given the evolutionary relationships for structurally large motifs (see below) in the MHC-II beta chain α-helix and the MHC-I heavy chain α-1 helix (Fig. 7).

**Figure 7.**
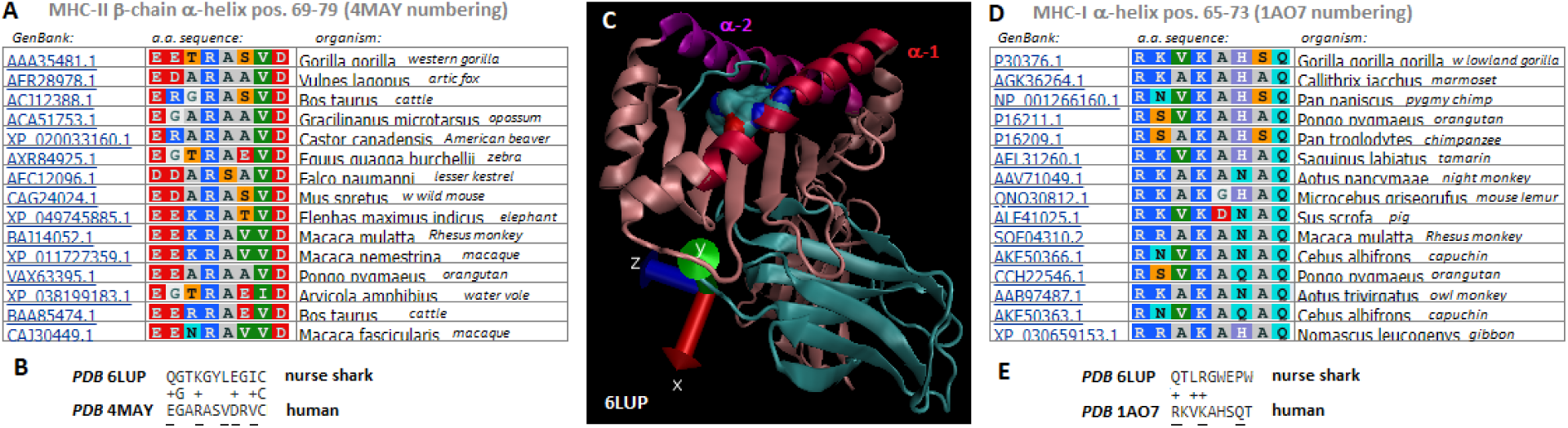
“*Pause”* motifs on MHC-II beta chain & MHC-I α-1 α-helices. (A) *NCBI Blast* results for 4MAY DQB1 (pos. 66-88) representative non-human hits in the top 100; zoomed to pos. 69-79. (B) Two sequence *Blast* of 4MAY DQB1 versus 6LUP MHC-I heavy chain. (C) Solved crystal structure of shark MHC-I molecule (*PDB* 6LUP; ref. 29). (D) *NCBI Blast* results for 1AO7 heavy chain (pos. 58-78) representative non-human hits in the top 100; zoomed to pos. 65-73. (E) Two sequence *Blast* 1AO7 versus 6LUP MHC-I heavy chain. Amino acids (a.a.) are shown in RASMOL colouring. *GenBank* accession numbers as annotated in *NCBI*, as are scientific species names; common names are abbreviated on same lines. MHC-I helices are colour-coded as in previous figures. The tandem tryptophans of the shark α-1 helix (also in E) are shown as space-filled by atom type with *VMD* (C). The light chain is show in *cyan*, heavy chain in *pink* with helices in separate colours: *purple* and *magenta* (α-2 helix), and *red* (α-1 helix). Highly conserved a.a. of the motifs are underlined in B & E, and non-matching a.a. of similar type are denoted by a ‘+’ symbol.

As recently described by Wu, Y., et al. (ref. 29), from bioinformatics and the crystallographic structure of the *nurse shark* MHC-I molecule (*PDB* 6LUP) it may be that an MHC-II like molecule came before MHC-I. Shown in Figure 7C is the shark MHC-I, and it is observed that the heavy-chain α-2 helix resembles the split α-helix of human MHC-II beta chains, and not the MHC-I α-2 helix of higher species [22, 23, 29]. A conserved motif is appreciated by *Blast* analysis (Fig. 7A), where is shown representative non-human species MHC-II beta chain α-helix sequences aligned in *COBALT*. It is notable the motif, E69-X-X-R72-X-X-V75-D76-X-V78 is present across widely separated species [30]. Indeed, the ‘break’ position, a.a. position 74, is mostly conserved (S/A/T/V), although the effect of E74 (*zebra*, *vole*, *cattle*) on the overall structure is conceivably disruptive (data not shown). Similarly, the tripartite combination of R65-X-X-K68-X-X-X-Q72, within the α-helix suggests a “bowl” along the α-1 helix of MHC-I that might also be an obstacle for (in this case) Vβ sliding down the IP. Note that this motif is also widely found across evolutionary time (Fig. 7D). Thus, Vα positions to a common site (as analysed) on most MHC-II beta chain α-helices, while Vβ is apparently free to assume variable positions along MHC-II alpha chain α-helices. Oppositely, Vα appears to be free to assume variable positions along the MHC-I α-2 helix, and this correlates with motifs of the two α-helices as indicated (Fig. 7). Interestingly, while Vβ is restricted to a common site along most MHC-I α-1 helices; this could perhaps be accomplished in a different fashion by the tandem bulky tryptophans in the shark, i.e., versus the R65/K68/Q72 *bowl* motif in higher species (Fig. 7C-7E).

## Discussion

Taken together, the suggestion that both Vα and Vβ began with restriction in terms of mobility along both MHC-I α-helices is interesting regarding this dual IP model for human TCR V-domains interacting with both MHC-I and MHC-II [25]. While the split into two helices was ostensibly lost for MHC-I α-2, i.e., allowing variable positioning of Vα, it was apparently not lost in the MHC-II beta lineage; thus, human Vα positioning is consistent between TCR restricted to MHC-II. Indeed, for the MHC-II restricted TCR, it is the Vβ domain which shows variability in positioning along the MHC-II alpha chain α-helix (Table 2). Clonotypic *dV*, ostensibly leading to a slide for any given V-domain of the TCR, is theoretically a product of the two CDR3-loops’ amino acid sequences [19]. Indeed, in the cases where V-domain mobility is limited, perhaps by the aforementioned motifs (Fig. 7), we have calculated that a potential variable encoded by the genetics of the V-domain is *time*. More specifically, the calculated sliding time for which the domain spends moving down its cognate α-helix (Table 2). This is a further indication that dynamics (i.e., protein domain motion per time) is proximal in pMHC discrimination by the TCR [5, 13, 18]. While H-bonding is envisioned to create a modeled frictional force (Fig. 1-Fig. 4), we have not investigated this here; rather we assume initial slip bonds allow the V-domain to move up the IP via the force (***F****_τ_*_∥_) and only re-forms H-bonding “friction” after the domain positions itself according to equalised acceleration up and down the IP. Clearly, H-bonding could modify acceleration, but obviously more sophisticated methods will be needed to make such estimates. Trajectories of V-domain dynamics are being targeted by techniques such as NMR [31], hydrogen-deuterium exchange MS [32], and other biochemical and biophysical approaches [16, 17, 33, 34]. It is also important to investigate TCR that bind pMHC in unusual orientations [35, 36]; which was not done, here.

Estimating the torque force (*τ*_θ_) resulting from a given V-domain rotating through the *dV*-specified *arc* has a component parallel force (***F****_τ_*_∥_) directed up the MHC α-helix (Fig. 1-4; Table 1). Crucially, the two IP of each MHC class-I and class-II protein face in *opposite* directions – from sharks to humans [22, 23, 29]. Thus, the specified parallel forces could reflect a mechanism present from near the beginning of the adaptive immune system, which includes a role for tangential shearing forces between the T-cell and APC membranes [13–17]. Clearly, such forces could orchestrate remarkable energy differences between target and ‘off-target’ TCR:pMHC complexes [37], facilitating the TCR *scanning* thousands of pMHC until finding the target ligand. The trajectory of each V-domain to the apex of each IP modeled here is suggested by a common, and opposite direction, yet there is clonotypic magnitude of ***F****_τ_*_∥_ values, i.e., due to variability in the location of the central cysteine relative to the pMHC groove (Table 1). While initial slip bonds could be envisioned to directly lead to variable *dx*/*dy* values estimated here for Vα on MHC-I α-2 helices and Vβ on MHC-II alpha α-helices, if a direct trajectory were in play (i.e., no sliding), it is curious that Vβ on MHC-I α-1 and Vα on MHC-II beta α-helices showed little variability in *dx*/*dy* values between different TCR (Table 2; Table 3 & **Supplement I**). Importantly, the physical chemistry of similar protein:protein binding, including with some *sIg*, does not indicate an enzymatic ‘lock-and-key’ mechanism [37–39]; nor, an ‘induced-fit’, per se [5, 38, 39]; rather, a common dynamic might be in play upon each pMHC encounter [11, 24, 25]. We can consider the initial binding position (which we can imagine) of a given V-domain (A) and its position at the apex of the MHC α-helix IP (B), which we can also imagine, as being in conformational equilibrium (ΔGc^‡^). As known by the *Curtin-Hammett principle*, the conformational equilibrium might not dictate product outcome (see below, ref. 40). Rather, it can be the ratio of the two transition states (ΔGa^‡^ and ΔGb^‡^) which decides a productive versus nonproductive TCR-pMHC encounter. However, much needs to be learned about the structure of water in the context of such a theory (ref. 42). Given *Brownian* motion [43, 44], thermal fluctuations (ref. 42), and fluid dynamics [45], it is perhaps premature to suggest even more than a simple theory for the mechanism of TCR antigen specificity..

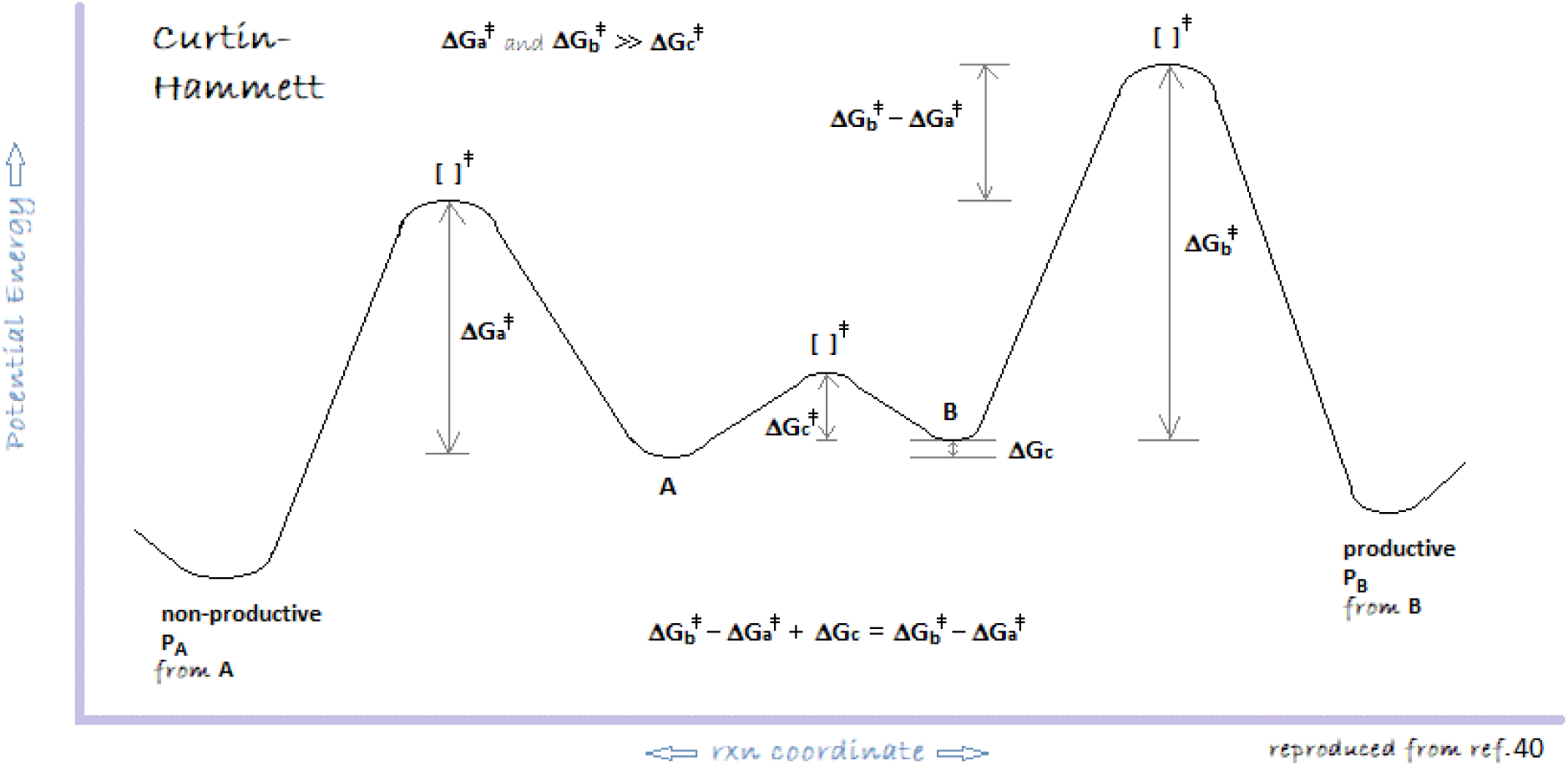

Finally, the T-cell clones from which DNA sequences of these TCR were identified, proteins isolated, and X-ray crystallography performed had already passed through rigorous TCR-selection in the foetal thymus [2, 12, 46, 47]. While the model might be viewed as highly speculative, there is striking structural linkage to it, as has been discussed. Clearly, it is interesting to consider that simply the timing of such *very* early dynamics may have begun to shape the biochemistry.

## Materials and Methods

*Availability of data and materials: PDB* files are public and available at the *NCBI* (http://www.ncbi.nlm.nih.gov); all analytics data are either in the paper, or supplementary materials; *Competing interests:* Dr. J. S. Murray and *Xenolaüs Genetics LLC* declare no competing interests regarding the research, or its publication; *Author Contributions:* J.S.M. did the research and wrote the manuscript.

### *PDB* file analysis

*VMD-1.9.1* software (http://www.ks.uiuc.edu) was used for analytical vector analysis of *PDB* files downloaded *via NCBI* (http://www.ncbi.nlm.nih.gov) from the *RSCB-PDB* (http://www.rscb.org); views normalized with the *VMD xyz-axis* tool; alpha carbon (Cα) main-chains in *new cartoon* were colour-coded in *VMD* to highlight specific protein regions; all alleles named per *NCBI* annotation. *Euler* methods (http://www.mathword.wolfram.com/EulerAngles. html) were the basis for specific angle analyses, as previously reported [11]. Briefly, three angles corresponding to the *twist*, *tilt*, and *sway* (orientation) of each domain over the pMHC were measured from fixed a.a. position Cα through the 20 structures (*n* = 40 separate V-domains; Tables 1-3; **Supplement I**; ref. 11, 48): (i) in-plane to the MHC-groove {*twist* = *ω*}, with (ii) displacements perpendicular to the groove {*tilt* = λ}, including (iii) side-to-side variation {*sway* = *σ*}. These measures were used to calculate the *cone shape* of each V-domain; specifically, the *ɸ*-angle and the cone radius (*r*) as described in **Results**. The other variable for cone-shape was the distance (*ρ*), which is the distance measured in *VMD* from the fixed a.a. position determining orientation and the V-domain central cysteine Cα (i.e., C21, C22, C23, depending on the structure).

### Calculation of the cone-slice volume element (*dV*)

Structural analysis included conversion of measurements in *VMD* into spherical coordinates used to solve individual triple-integral equations for all V-domains [11]. Briefly, the general equation for the volume element (*dV*) (with defined trigonometry) was used as previously described (ref. 11):

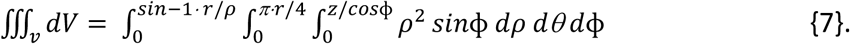

Specific *VMD* measurements from the indicated *PDB* files were the angle-*ɸ*: calculated by subtraction of the *tilt* (λ) angle from normal (180°) at the fixed a.a. position used to describe orientation relative to the pMHC groove for each V-domain; the *ɸ*-angle integrand is the first from the left in the general equation {7}. The second integrand is for *θ*_s_, which is the arc distance created by the ‘sweep’ of *dV*, i.e., *dθ*. Finally, the third integrand is for the distance (*ρ*), which is the measured distance from the same fixed a.a. position describing orientation to the V-domain’s central cysteine. Thus, only the angle-*ɸ* required conversion to a distance (multiplying by *π*/180) to obtain the Å^3^ volume for *dV*.

### Estimation of V-domain slope-parallel torque (*F_τ_*_∥_) from *dV* and the moment of inertia (*I*) of a flat-top cone

It followed that the general equation for the moment of inertia (***I***) of a flat-top cone (ref. 27) could be used to calculate torque from the equation:

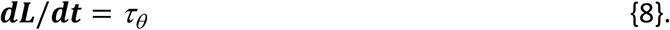

Where, momentum (***L***) is the cross-product of (***I***) and velocity through *dθ*, that is, the arc distance (*θ*_s_) traveled by the central cysteine in each *dV*. Thus, we use the first derivative of momentum to define overall torque (τ_θ_). As previously indicated, it was necessary to estimate direction of the overall torque vector from the crystallographic structure in order to determine the component of the vector running parallel (and up) the MHC α-helix inclined plane. In fact, while we measured most of these vectors to be opposite the ***F***w vector to the IP, and thus multiplication by the *sine* of angle-γ (or γ_2_) could derive the slope-parallel force (*Fτ* _∥_), in the case of the Vβ domain of MHC-I restricted TCR, the overall torque was a slope-perpendicular force vector (can be seen in Fig. 2 & 4); in these cases, we could divide by the *cosine* of the angle-γ to obtain ***F****_τ_*_∥_. Individual V-domain solutions to the series of equations are shown in the diagrams of Figure 1 through Figure 4.

### Estimation of V-domain slide time (*t*) on MHC α-helix IP

It follows from the static position of the V-domain as it is bound to the IP of any given MHC α-helix, that acceleration up and down the IP must be equivalent [26, 27]. Thus, the force vector counteracting the slope-parallel component of torque involves ‘sliding’ down the IP. While we use the ground-state coordinates to estimate these torque values, it does not necessarily follow that the ground-state coordinates ‘define’ this initial position. It is rather, an estimate using the ground-state coordinates. The other crucial assumption here, which is consistent with using velocity through *dθ* to determine torque, is that between structures the arc-distance traveled takes precisely 1.00 *μ*s. This is an assumption needed to isolate the torque, and indeed, acceleration vectors. Again, it is an estimate. The derivation first uses the law of energy conservation to obtain the kinetic energy of the V-domain at the ground-state CDR2 contact along the IP, i.e., the point after sliding (***a***). See equations leading to equation {3}:

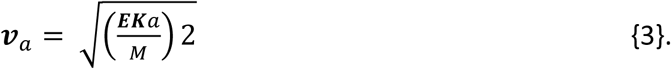

This is *Newton’s kinetic energy* equation solving for velocity and using the ***EK****_a_* value previously obtained from the *first law of thermodynamics*. Once velocity at position (***a***) is known, the *third kinematic equation* {4} was used to calculate acceleration down the IP at position (***a***) [26, 27].

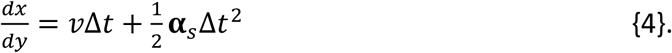

The use of the equation requires the distance *dx*/*dy*, and here the value estimated from the ground-state position of the central cysteine relative to the apex of the IP, i.e., angle-*η* and the distance (*ψ*) (Fig. 5 & 6), was used. Finally, once acceleration was estimated, this value (expressed as Å/*μ*s/***t***) would need to be equal to acceleration from the slope-parallel component of torque, i.e., for the V-domain to obtain a stationary ground-state structure. See equations, {5} and {6}.

### Contacts analysis

All measures were performed with *VMD-1.9.1* as specified in **Results**. Closest contacts in *angstroms* (Å) were determined by the *bond-label* tool, examining appropriate coordinates between structures. Individual atomic contacts are named per software annotation. These were converted to single letter a.a. code in the text.

### Bioinformatics analysis of MHC α-helix motifs

All bioinformatics data (*PDB* files, sequences, *COBALT* multiple alignments, *BLAST* searches) were performed with algorithms available at the *NCBI* (http://www.ncbi.nlm.nih.gov). Briefly, a.a. sequences for MHC protein chains for the full length of the inclined plane α-helix were imported from the TCR:pMHC structure file into the *BLAST* function. *BLAST* was with the *blastp* (protein-protein) algorithm. *Homo sapiens* were excluded with the ‘*search-set*’ tool. Parameters were set to 100 hits, 200,000 threshold, word-size, 2; matrix, PAM30 with Gap Costs set to existence: 9, and extension: 1. Representative species MHC matches were chosen for multiple alignment, and a.a. shown in RASMOL colouring using the ‘*graph overview*’ tool to zoom into the sequence of interest. The original outputs were incorporated into Fig. 7 with original annotation and common names included from *NCBI taxa* files.

## Acknowledgements

Public access to crystallographic coordinates, genetic sequences and computer software made this work possible. Thanks to E.H. Murray, M.D., for reviewing the manuscript.

## Appendix

Supplementary data (**Supplement I**) are available in the on-line version: “C:\Users\xenol\OneDrive\Desktop\TCR\Supplement I.pdf”

## Notes

### Competing Interest Statement

The authors have declared no competing interest.

### Summary of Updates

Considerable changes to text with additional references.

