## Supplement I for "Dichotomy in TCR V-domain dynamics binding the opposed inclined planes of pMHC-II and pMHC-I α-helices"

### A. Linear regression analysis of V-domains

| | | | | | |
|---|---|---|---|---|---|
| 2IAM/Vα | 1702.87 | 27.76 | 204.3 | 0.136 | 15.25 |
| 4MAY/Vα | 615.02 | 26.72 | 125.12 | 0.214 | 14.76 |
| 1J8H/Vα | 746.24 | 26.46 | 125.33 | 0.207 | 14.55 |
| 4E41/Vα | 1512.33 | 19.68 | 188.8 | 0.104 | 10.99 |
| 4OZG/Vα | 1657.12 | 27.32 | 200.75 | 0.136 | 15.01 |
| 1FYT/Vα | 826.36 | 26.38 | 137.28 | 0.192 | 14.52 |
| 4H1L/Vα | 1781.86 | 28.64 | 196.54 | 0.146 | 15.66 |
| 6CQR/Vα | 718.16 | 19.04 | 129.67 | 0.147 | 10.67 |
| 4OZH/Vα | 2029.75 | 27.14 | 220.95 | 0.123 | 14.91 |
| 4OZF/Vα | 1395.98 | 26.1 | 173.04 | 0.151 | 14.34 |

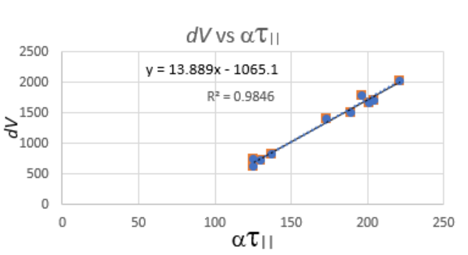


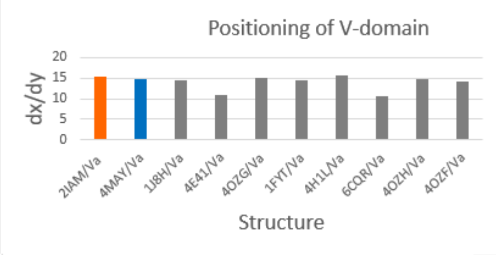


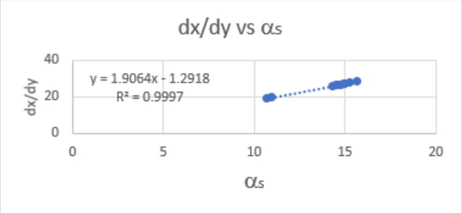


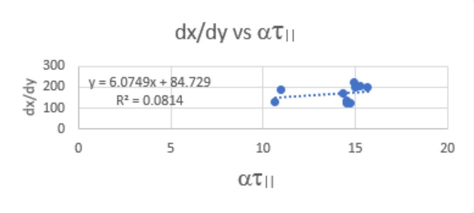


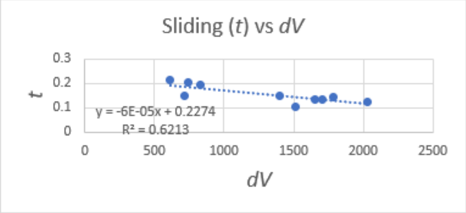


| | | | | | |
|---|---|---|---|---|---|
| 2IAM/Vβ | 296.64 | 46.17 | 0.266 | 12.29 | 6.82 |
| 4MAY/Vβ | 571.39 | 82.65 | 0.377 | 31.14 | 16.79 |
| 1J8H/Vβ | 1734.3 | 107.66 | 0.255 | 27.47 | 14.42 |
| 4E41/Vβ | 405.77 | 49.75 | 0.239 | 11.87 | 6.38 |
| 4OZG/Vβ | 276.3 | 42.16 | 0.54 | 22.76 | 12.26 |
| 1FYT/Vβ | 1746.97 | 98.75 | 0.268 | 26.43 | 14.13 |
| 4H1L/Vβ | 1198.48 | 83.71 | 0.102 | 8.57 | 4.82 |
| 6CQR/Vβ | 745.55 | 70.27 | 0.38 | 26.71 | 14.31 |
| 4OZH/Vβ | 266.96 | 39.63 | 0.584 | 23.13 | 12.44 |
| 4OZF/Vβ | 553.87 | 59.28 | 0.46 | 27.26 | 14.58 |

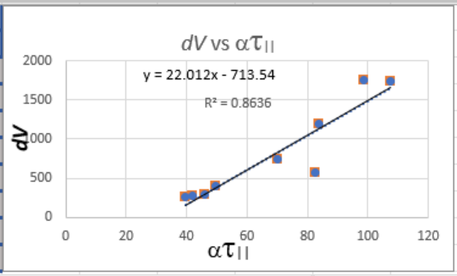


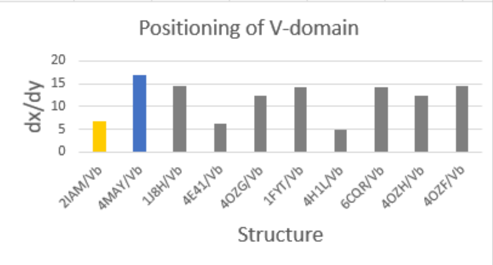


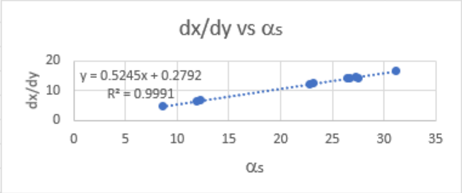


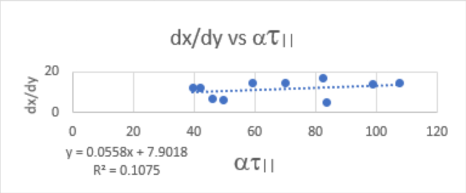


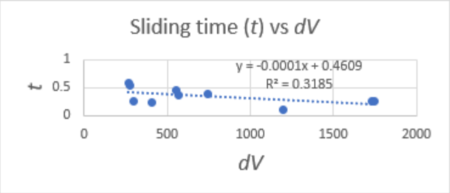

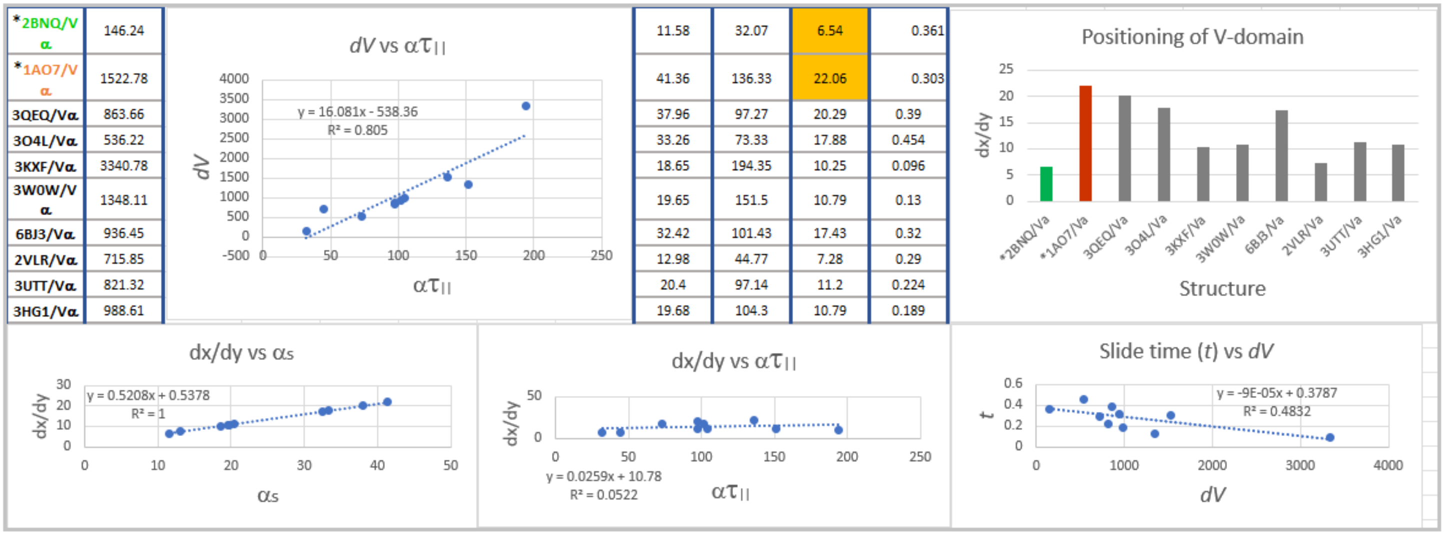

| | | | | | |
|---|---|---|---|---|---|
| *2BNQ/Vα | 146.24 | 11.58 | 32.07 | 6.54 | 0.361 |
| *1AO7/Vα | 1522.78 | 41.36 | 136.33 | 22.06 | 0.303 |
| 3QEQ/Vα | 863.66 | 37.96 | 97.27 | 20.29 | 0.39 |
| 3O4L/Vα | 536.22 | 33.26 | 73.33 | 17.88 | 0.454 |
| 3KXF/Vα | 3340.78 | 18.65 | 194.35 | 10.25 | 0.096 |
| 3W0W/Vα | 1348.11 | 19.65 | 151.5 | 10.79 | 0.13 |
| 6BJ3/Vα | 936.45 | 32.42 | 101.43 | 17.43 | 0.32 |
| 2VLR/Vα | 715.85 | 12.98 | 44.77 | 7.28 | 0.29 |
| 3UTT/Vα | 821.32 | 20.4 | 97.14 | 11.2 | 0.224 |
| 3HG1/Vα | 988.61 | 19.68 | 104.3 | 10.79 | 0.189 |



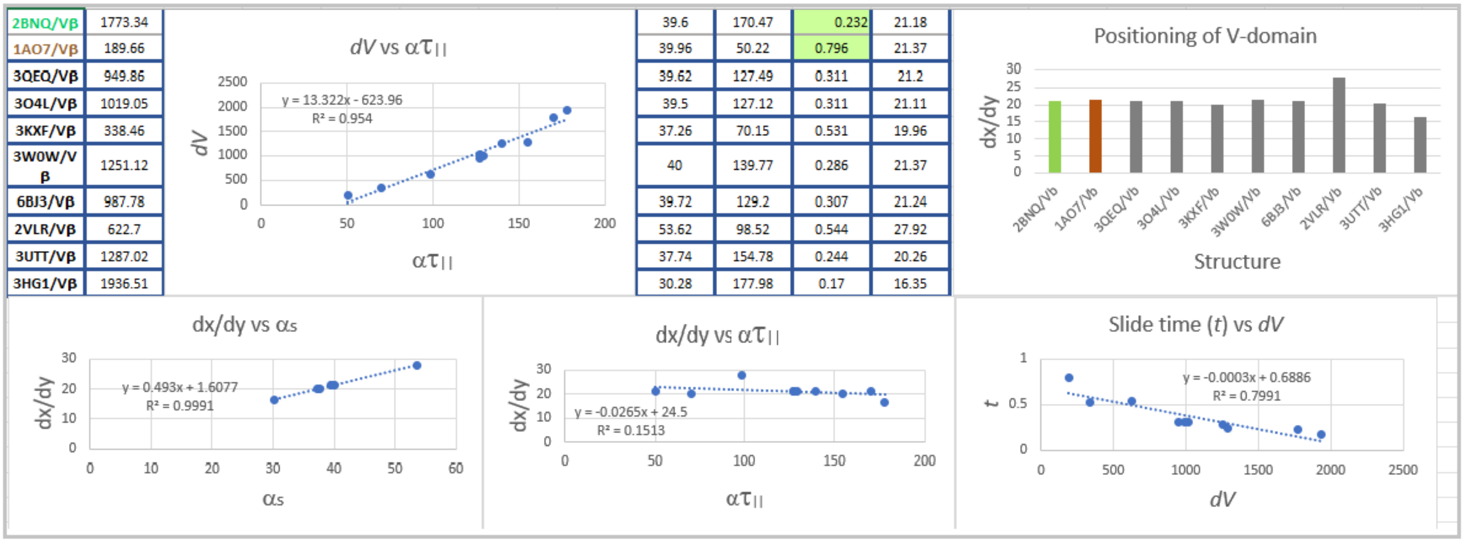

| | | | | | |
|---|---|---|---|---|---|
| 2BNQ/Vβ | 1773.34 | 39.6 | 170.47 | 0.232 | 21.18 |
| 1AO7/Vβ | 189.66 | 39.96 | 50.22 | 0.796 | 21.37 |
| 3QEQ/Vβ | 949.86 | 39.62 | 127.49 | 0.311 | 21.2 |
| 3O4L/Vβ | 1019.05 | 39.5 | 127.12 | 0.311 | 21.11 |
| 3KXF/Vβ | 338.46 | 37.26 | 70.15 | 0.531 | 19.96 |
| 3W0W/Vβ | 1251.12 | 40 | 139.77 | 0.286 | 21.37 |
| 6BJ3/Vβ | 987.78 | 39.72 | 129.2 | 0.307 | 21.24 |
| 2VLR/Vβ | 622.7 | 53.62 | 98.52 | 0.544 | 27.92 |
| 3UTT/Vβ | 1287.02 | 37.74 | 154.78 | 0.244 | 20.26 |
| 3HG1/Vβ | 1936.51 | 30.28 | 177.98 | 0.17 | 16.35 |



**B. Series of equations solutions for estimation of V-domain ‘slide times’ ($t$) on MHC α-helix IP based on energy conservation (equal acceleration up and down IP) at the ground-state location of CDR2 contact/central cysteine position.**

### 2IAM-Vα

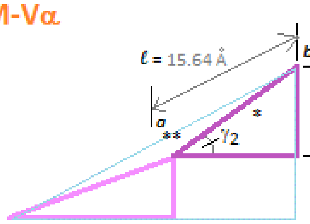


$htb = \sin\gamma_2\,(\ell)$
0.624(15.64)
=9.76 Å

$^*dx/dy = \sin\eta\,(\psi)$
$\sin(46.89)(20.89)$
=15.25 Å

$^{**}hta = \sin\gamma_2\,(\ell - dx/dy)$
0.624(0.39)
=0.243 Å

$EKb = 0$
$EPb = Mg\,(htb)$
= 1.19 kDa · Å/µs$^2$
x 9.76 Å
= 11.64 kDa · Å$^2$/µs$^2$

$M$ = 12.17 kDa
$g$ = 0.098 Å/µs$^2$
$\gamma_2$ = 38.61°

$EPa = Mg\,(hta)$
= 1.19 kDa · Å/µs$^2$
x 0.243 Å
= 0.290 kDa · Å$^2$/µs$^2$

$EKa = EPb - EPa$
= 11.64 - 0.290
= 11.35 kDa · Å$^2$/µs$^2$

where,
$EKa = 1/2\,M\,v^2$ = 11.35 kDa · Å$^2$/µs$^2$
$v^2$ = 2(11.35)/12.17
{3} $v_a = \sqrt{1.87}$ = 1.37 Å/µs

∴
{4} $dx/dy = v\,\Delta t + 1/2\,\alpha_s\,\Delta t^2$
$\alpha_s = \dfrac{15.25\text{Å} - 1.37\text{Å}\ \Delta t}{\Delta t^2\ \ \mu s} \times 2$
$\alpha_s$ = 27.76 Å/µs/$t$

{1} $F\tau_{||}$ = 3984.35 kDa · Å/µs$^2$ ($\sin\gamma_2$)
= 2486.30 kDa · Å/µs$^2$

where,
$F = M\,\alpha\,\tau_{||}$
$\alpha\,\tau_{||} = \dfrac{2486.30\ \text{kDa} \cdot \text{Å}/\mu s^2}{12.17\ \text{kDa}}$
= 204.30 Å/µs$^2$

∴
{5} $\alpha_s = \alpha\,\tau_{||}$
27.76 Å/µs/$t$ = 204.30 Å/µs$^2$
{6} $t = \dfrac{27.76\ \text{Å}\ \ \mu s^2}{\mu s\ 204.30\ \text{Å}}$
= 0.136 µs "slide time"

### 2IAM-Vβ

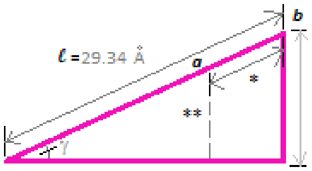


$htb = \sin\gamma\,(\ell)$
0.343 (29.34)
=10.05 Å

$^*dx/dy = \sin\eta\,(\psi)$
$\sin(19.12)(20.82)$
= 6.82 Å

$^{**}hta = \sin\gamma\,(\ell - dx/dy)$
0.343 (22.52)
= 7.72 Å

$EKb = 0$
$EPb = Mg\,(htb)$
= 1.20 kDa · Å/µs$^2$
x 10.05 Å
= 12.10 kDa · Å$^2$/µs$^2$

$M$ = 12.29 kDa
$g$ = 0.098 Å/µs$^2$
$\gamma$ = 20.04°

$EPa = Mg\,(hta)$
= 1.20 kDa · Å/µs$^2$
x 7.72 Å
= 9.30 kDa · Å$^2$/µs$^2$

$EKa = EPb - EPa$
= 12.10 - 9.30
= 2.80 kDa · Å$^2$/µs$^2$

where,
$EKa = 1/2\,M\,v^2$ = 2.80 kDa · Å$^2$/µs$^2$
$v^2$ = 2( 2.80 )/ 12.29
{3} $v = \sqrt{0.456}$ = 0.675 Å/µs

∴
{4} $dx/dy = v\,\Delta t + 1/2\,\alpha_s\,\Delta t^2$
$\alpha_s = \dfrac{6.82\ \text{Å} - 0.675\text{Å}\ \Delta t}{\Delta t^2\ \ \mu s} \times 2$
$\alpha_s$ = 12.29 Å/µs/$t$

{1} $F\tau_{||}$ = 1655.94 kDa · Å/µs$^2$ ($\sin\gamma$)
= 567.45 kDa · Å/µs$^2$

where,
$F = M\,\alpha\,\tau_{||}$
$\alpha\,\tau_{||} = \dfrac{567.45\ \text{kDa} \cdot \text{Å}/\mu s^2}{12.29\ \text{kDa}}$
= 46.17 Å/µs$^2$

∴
{5} $\alpha_s = \alpha\,\tau_{||}$
12.29 Å/µs/$t$ = 46.17 Å/µs$^2$
{6} $t = \dfrac{12.29\ \text{Å}\ \ \mu s^2}{\mu s\ 46.17\ \text{Å}}$
= 0.266 µs "slide time"

### 4MAY-Vα

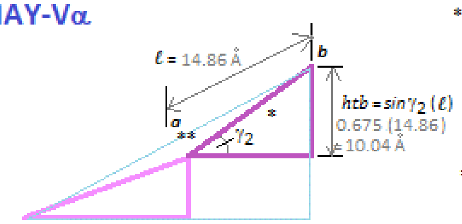


$^{*}dx/dy = \sin\eta\,(\psi)$
$\sin(31.10)(28.58)$
$= 14.76$ Å

$^{**}hta = \sin\gamma_2\,(\ell - dx/dy)$
$0.675\,(14.86 - 14.76)$
$= 0.082$ Å

$EKb = 0$
$EPb = Mg\,(htb)$
$= 1.20$ kDa · Å/µs$^2$
x 10.04 Å
$= 12.03$ kDa · Å$^2$/µs$^2$

$M = 12.23$ kDa
$g = 0.098$ Å/µs$^2$
$\gamma_2 = 42.48°$

$EPa = Mg\,(hta)$
$= 1.20$ kDa · Å/µs$^2$
x 0.068 Å
$= 0.082$ kDa · Å$^2$/µs$^2$

$EKa = EPb - EPa$
$= 12.03 - 0.082$
$= 11.95$ kDa · Å$^2$/µs$^2$

where,
$EKa = 1/2\,M v^2 = 11.95$ kDa · Å$^2$/µs$^2$
$v^2 = 2(11.95)/12.23$
{3} $v = \sqrt{1.95} = 1.40$ Å/µs

{4} ∴ $dx/dy = v\,\Delta t + 1/2\,\alpha_s\,\Delta t^2$
$\alpha_s = \dfrac{14.76\text{ Å} - 1.40\text{ Å}\ \cancel{\Delta t}}{\Delta t^{\cancel{2}}\ \mu s} \times 2$
$\alpha_s = 26.72$ Å/µs/$t$

{1} $F\tau_{||} = 2265.63$ kDa · Å/µs$^2$ ($\sin\gamma_2$)
$= 1530.05$ kDa · Å/µs$^2$

where,
$F = M\,\alpha\,\tau_{||}$
$\alpha\,\tau_{||} = \dfrac{1530.05\ \cancel{kDa} \cdot \text{Å}/\mu s^2}{12.23\ \cancel{kDa}}$
$= 125.12$ Å/µs$^2$

{5} ∴ $\alpha_s = \alpha\,\tau_{||}$
$26.72$ Å/µs/$t$ $= 125.12$ Å/µs$^2$
{6} $t = \dfrac{26.72\ \cancel{Å}\ \mu s^{\cancel{2}}}{\cancel{\mu s}\ 125.12\ \cancel{Å}}$
$= 0.214$ µs "slide time"

### 4MAY-Vβ

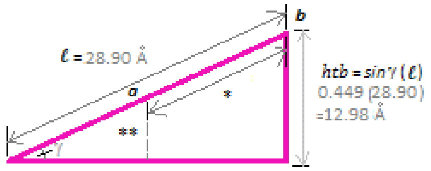


$^{*}dx/dy = \sin\eta\,(\psi)$
$\sin(43.89)(24.22)$
$= 16.79$ Å

$^{**}hta = \sin\gamma\,(\ell - dx/dy)$
$0.449\,(12.11)$
$= 5.44$ Å

$EKb = 0$
$EPb = Mg\,(htb)$
$= 1.16$ kDa · Å/µs$^2$
x 12.98 Å
$= 15.10$ kDa · Å$^2$/µs$^2$

$M = 11.87$ kDa
$g = 0.098$ Å/µs$^2$
$\gamma = 26.69°$

$EPa = Mg\,(hta)$
$= 1.16$ kDa · Å/µs$^2$
x 5.44 Å
$= 6.33$ kDa · Å$^2$/µs$^2$

$EKa = EPb - EPa$
$= 15.10 - 6.33$
$= 8.77$ kDa · Å$^2$/µs$^2$

where,
$EKa = 1/2\,M v^2 = 8.77$ kDa · Å$^2$/µs$^2$
$v^2 = 2(8.77)/11.87$
{3} $v = \sqrt{1.48} = 1.22$ Å/µs

{4} ∴ $dx/dy = v\,\Delta t + 1/2\,\alpha_s\,\Delta t^2$
$\alpha_s = \dfrac{16.79\text{ Å} - 1.22\text{ Å}\ \cancel{\Delta t}}{\Delta t^{\cancel{2}}\ \mu s} \times 2$
$\alpha_s = 31.14$ Å/µs/$t$

{1} $F\tau_{||} = 2184.14$ kDa · Å/µs$^2$ ($\sin\gamma$)
$= 981.04$ kDa · Å/µs$^2$

where,
$F = M\,\alpha\,\tau_{||}$
$\alpha\,\tau_{||} = \dfrac{981.04\ \cancel{kDa} \cdot \text{Å}/\mu s^2}{11.87\ \cancel{kDa}}$
$= 82.65$ Å/µs$^2$

{5} ∴ $\alpha_s = \alpha\,\tau_{||}$
$31.14$ Å/µs/$t$ $= 82.65$ Å/µs$^2$
{6} $t = \dfrac{31.14\ \cancel{Å}\ \mu s^{\cancel{2}}}{\cancel{\mu s}\ 82.65\ \cancel{Å}}$
$= 0.377$ µs "slide time"

### 2BNQ-Vα

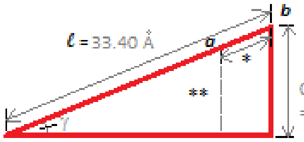


$htb = \sin\gamma\,(\ell)$
0.436 (33.40)
= 14.57 Å

$^{*}dx/dy = \sin\eta\,(\psi)$
sin (16.30)(23.31)
= 6.54 Å

$^{**}hta = \sin\gamma\,(\ell - dx/dy)$
0.436 (26.86)
= 11.72 Å

$EKb = 0$
$EPb = Mg\,(htb)$
= 1.15 kDa · Å/µs$^2$
x 14.57 Å
= 16.76 kDa · Å$^2$/µs$^2$

$M$ = 11.73 kDa
$g$ = 0.098 Å/µs$^2$
$\gamma$ = 25.86°

$EPa = Mg\,(hta)$
= 1.15 kDa · Å/µs$^2$
x 11.72 Å
= 13.48 kDa · Å$^2$/µs$^2$

$EKa = EPb - EPa$
= 16.76 - 13.48
= 3.28 kDa · Å$^2$/µs$^2$

where,
$EKa = 1/2\,M\,v^2$ = 3.28 kDa · Å$^2$/µs$^2$
$v^2$ = 2( 3.28 )/ 11.73
{3} $v_a = \sqrt{0.560}$ = 0.748 Å / µs

∴
{4} $dx/dy = v\,\Delta t + 1/2\,\alpha_S\,\Delta t^2$

$\alpha_S$ = (6.54 Å - 0.748 Å ~~Δt~~ / Δ t ~~2~~ µs) x 2

$\alpha_S$ = 11.58 Å / µs / $t$

{1} $F\tau_{||}$ = 862.34 kDa · Å/µs$^2$ (sin γ)
= 376.13 kDa · Å/µs$^2$

where,
$F = M\,\alpha\,\tau_{||}$

$\alpha\,\tau_{||}$ = 376.13 ~~kDa~~ · Å/µs$^2$ / 11.73 ~~kDa~~

= 32.07 Å/µs$^2$

∴
{5} $\alpha_S = \alpha\,\tau_{||}$

11.58 Å / µs / $t$ = 32.07 Å/µs$^2$

{6} $t$ = 11.58 ~~Å~~ µs~~2~~ / ~~µs~~ 32.07 ~~Å~~

= 0.361 µs "slide time"

### 2BNQ-Vβ

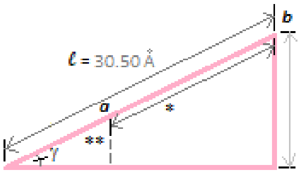


$htb = \sin\gamma\,(\ell)$
0.456(30.50)
=13.92Å

$^{*}dx/dy = \sin\eta\,(\psi)$
sin (46.28)(29.31)
= 21.18 Å

$^{**}hta = \sin\gamma\,(\ell - dx/dy)$
0.456 (9.32)
= 4.25 Å

$EKb = 0$
$EPb = Mg\,(htb)$
= 1.18 kDa · Å/µs$^2$
x 13.92 Å
= 16.48 kDa · Å$^2$/µs$^2$

$M$ = 12.08 kDa
$g$ = 0.098 Å/µs$^2$
$\gamma$ = 27.16°

$EPa = Mg\,(hta)$
= 1.18 kDa · Å/µs$^2$
x 4.25 Å
= 5.03 kDa · Å$^2$/µs$^2$

$EKa = EPb - EPa$
= 16.48 - 5.03
= 11.45 kDa · Å$^2$/µs$^2$

where,
$EKa = 1/2\,M\,v^2$ = 11.45 kDa · Å$^2$/µs$^2$
$v^2$ = 2(11.45 )/ 12.08
{3} $v = \sqrt{1.90}$ = 1.38 Å / µs

∴
{4} $dx/dy = v\,\Delta t + 1/2\,\alpha_S\,\Delta t^2$

$\alpha_S$ = (21.18 Å -1.38 Å ~~Δt~~ / Δ t ~~2~~ µs) x 2

$\alpha_S$ = 39.60 Å / µs / $t$

{2} $F\tau_{||}$ = 4019.13 kDa · Å/µs$^2$ ( $\frac{\sin\gamma}{\cos\gamma}$ )
= 2062.01 kDa · Å/µs$^2$

where,
$F = M\,\alpha\,\tau_{||}$

$\alpha\,\tau_{||}$ = 2062.01 ~~kDa~~ · Å/µs$^2$ / 12.08 ~~kDa~~

= 170.47 Å/µs$^2$

∴
{5} $\alpha_S = \alpha\,\tau_{||}$

39.60 Å / µs / $t$ = 170.47 Å/µs$^2$

{6} $t$ = 39.60 ~~Å~~ µs~~2~~ / ~~µs~~ 170.47 ~~Å~~

= 0.232 µs "slide time"

## 1AO7-Vα

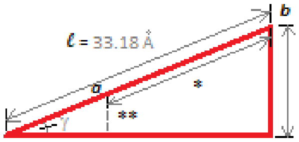


htb = sinγ (ℓ)
0.443 (33.18)
=14.71Å

*dx/dy = sin η (ψ)
sin (55.55)(26.75)
=22.06 Å

**hta = sinγ (ℓ - dx/dy)
0.443 (11.12)
= 4.93 Å

EKb = 0
EPb = Mg (htb)
=1.18 kDa · Å/µs2
x 14.71 Å
=17.28 kDa · Å2/ µs2

M = 11.99 kDa
g = 0.098 Å/µs2
γ = 26.32 °

EPa = Mg (hta)
=1.18 kDa · Å/µs2
x 4.93 Å
= 5.79 kDa · Å2/ µs2

EKa = EPb - EPa
=17.28 - 5.79
= 11.49 kDa · Å2/ µs2

where,
$EKa = 1/2\, M v^2 = 11.49$ kDa · Å2/ µs2
$v^2 = 2(11.49)/11.99$
{3} $v_a = \sqrt{1.92} = 1.38$ Å / µs

∴
{4} $dx/dy = v\,\Delta t + 1/2\, \alpha_S\, \Delta t^2$
$\alpha_S = \dfrac{22.06\ \text{Å} - 1.38\ \text{Å}\ \Delta t}{\Delta t^2\ \ \mu s} \times 2$
$\alpha_S = 41.36$ Å / µs / t

{1} $F\tau_{||} = 3686.70$ kDa · Å/µs2 (sinγ)
= 1634.62 kDa · Å/µs2

where,
$F = M\, \alpha\, \tau_{||}$
$\alpha\tau_{||} = \dfrac{1634.62\ \text{kDa} \cdot \text{Å/µs}^2}{11.99\ \text{kDa}}$
= 136.33 Å/µs2

∴
{5} $\alpha_S = \alpha\, \tau_{||}$
41.36Å / µs / t = 136.33Å/µs2
{6} $t = \dfrac{41.36\text{Å}\ \ \mu s^2}{\mu s\ \ 136.33\text{Å}}$
= 0.303 µs "slide time"

## 1AO7-Vβ

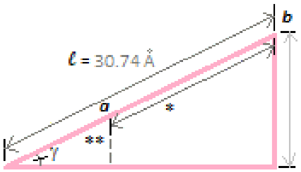


htb = sinγ (ℓ)
0.460 (30.74)
=14.13Å

*dx/dy = sin η (ψ)
sin (42.13)(31.86)
=21.37 Å

**hta = sinγ (ℓ - dx/dy)
0.460 (9.37)
= 4.31 Å

EKb = 0
EPb = Mg (htb)
= 1.18 kDa · Å/µs2
x 14.13 Å
=16.62 kDa · Å2/ µs2

M = 11.99 kDa
g = 0.098 Å/µs2
γ = 27.36°

EPa = Mg (hta)
=1.18 kDa · Å/µs2
x 4.31 Å
= 5.06 kDa · Å2/ µs2

EKa = EPb - EPa
= 16.62 - 5.06
= 11.56 kDa · Å2/ µs2

where,
$EKa = 1/2\, M v^2 = 11.56$ kDa · Å2/ µs2
$v^2 = 2(11.56)/11.99$
{3} $v_a = \sqrt{1.93} = 1.39$ Å / µs

∴
{4} $dx/dy = v\,\Delta t + 1/2\, \alpha_S\, \Delta t^2$
$\alpha_S = \dfrac{21.37\ \text{Å} - 1.39\ \text{Å}\ \Delta t}{\Delta t^2\ \ \mu s} \times 2$
$\alpha_S = 39.96$ Å / µs / t

{2} $F\tau_{||} = 1162.29$ kDa · Å/µs2 $\left(\dfrac{\sin\gamma}{\cos\gamma}\right)$
= 602.09 kDa · Å/µs2

where,
$F = M\, \alpha\, \tau_{||}$
$\alpha\tau_{||} = \dfrac{602.09\ \text{kDa} \cdot \text{Å/µs}^2}{11.99\ \text{kDa}}$
= 50.22 Å/µs2

∴
{5} $\alpha_S = \alpha\, \tau_{||}$
Å / µs / t = 39.96 Å/µs2
{6} $t = \dfrac{39.96\ \text{Å}\ \ \mu s^2}{\mu s\ \ 50.22\ \text{Å}}$
= 0.796 µs "slide time"

#### C. Broad analysis of relative CDR2-contacts along MHC α-helices in solved TCR:pMHC structures.

**Table 3A. Relative mobility of CDR2-loop along MHC α-helices in solved TCR:pMHC#**

MHC-I α-helix location α-helix location

| ***PDB*/Vα** (MHC allele) | Top α-2 (151-161) | Middle α-2 (162-167) | Bottom α-2 (168-173) | ***PDB*/Vβ** | Top α-1 (58-64) | Middle α-1 (65-72) | Bottom α-1 73-78) |
|---|---|---|---|---|---|---|---|
| 1AO7 (A*0201) | | N52:ND2-E166:OE2 **3.12** | | 1AO7 | | I54:CD1-Q72:CG **6.52** | |
| 2BNQ (A*0201) | S53:OG-Q155NE2 **2.75** | | | 2BNQ | | A51:N-Q72:OE1 **2.93** | |
| 3HG1 (A*0201) | | N53:ND2-E166:OE1 **6.62** | | 3HG1 | | Q55:OE1-Q72:NE2 **3.98** | |
| 3O4L (A*0201) | | S52:OG-E166:OE2 **5.08** | | 3O4L | | N52:ND2-Q72:OE1 **4.32** | |
| 3QEQ (A*0201) | K50:CB-A158:CB **4.43** | | | 3QEQ | | D54:OD2-Q72:NE2 **3.09** | |
| 3W0W (A*2402) | S52:OG-E154:OE2 **2.25** | | | 3W0W | | V52:O-Q72:NE2 2.99 | |
| 3KXF (B*3508) | F53:CD1-R157:CD **3.43** | | | 3KXF | | E52:OE1-Q72:NE2 **10.97** | |
| 6BJ3 (B*3501) | | D54:OD2-L163:N **3.02** | | 6BJ3 | | S51:OG-Q72:NE2 **3.95** | |
| 2VLR (A*0201) | S31:OG-Q155:N **3.75** | | | 2VLR | | | I53:CD-V76:CG2 **3.94** |
| 3UTT (A*0201) | Y51:CD1-A158:CB **7.60** | | | 3UTT | | V53:CG2-Q72:CD **3.66** | |

#CDR2-loop contact, in Å, as measured with *VMD* (http://www.ks.uiuc.edu). Colour code: *red*: 2.0 to 2.5 H-bond; *purple*: 2.6 to 3. 5 H-bond; *green*: 3.6 and above; *blue*: hydrophobic. Note: this is an a.a. of CDR2 that is near the apex of the loop, in some cases closer contacts can be found at the bases of the loop; however, these contacts do not correlate with the location of the central cysteine; thus, relative position of the domain along the α-helix IP.

**Table 3B. Relative mobility of CDR2-loop along MHC α-helices in solved TCR:pMHC[#]**

| MHC-II | α-helix location | | | | α-helix location | | |
|---|---|---|---|---|---|---|---|
| MHC-II ***PDB*/Vα** **(MHC allele)** | Top DRB/DQB (66-74) | Middle DRB/DQB (75-81) | Bottom DRB/DQB (82-88) | ***PDB*/Vβ** **(MHC allele)** | Top DRA/DQA (57-64) | Middle DRA/DQA (65-70) | Bottom DRA/DQA (71-76) |
| 2IAM (DRB1) | | S50:OG-D76:OD2 **3.77** | | 2IAM (DRA) | I54:CD1-A61:CB 4.40 | | |
| 4MAY (DQB1) | | N53:OD1-R77NH2 **7.61** | | 4MAY (DQA1) | | T50:OG1-H68:ND1 **3.27** | |
| 4E41 (DRB1) | | S50:OG-D76:OD2 **5.37** | | 4E41 (DRA) | R53:NH1-L60:O **4.15** | | |
| 1J8H (DRB1) | | S51:OG-T77OG1 **5.66** | | 1J8H (DRA) | | D51:OD1-K67:NZ **2.54** | |
| 4H1L (DRB3) | | F50:O-N77:ND2 **3.81** | | 4H1L (DRA) | I49:CD1-L60:CB **3.30** | | |
| 6CQR (DRB1) | | L58:CG-T77:CG2 **4.35** | | 6CQR (DRA) | | E60:OE2-K67:NZ **2.83** | |
| 4OZH (DQB1) | | K58:NZ-D76:OD2 **4.83** | | 4OZH (DQA1) | Q57:CG-A61:CB **3.85** | | |
| 4OZF (DQB1) | | K58:NZ-D76:OD2 **4.17** | | 4OZF (DQA1) | | N63:OD1-K67:NZ **3.50** | |
| 4OZG (DQB1) | | L57:CD2-R77:CG **4.03** | | 4OZG | | Q57:CD-V65:CG2 **3.94** | |
| 1FYT (DRB1) | | S51:CB-T77:CG2 **4.62** | | 1FYT | | D51:OD1-K67:NZ **2.67** | |

[#]CDR2-loop contact, in Å, as measured with *VMD* (http://www.ks.uiuc.edu). Colour code: *red*: 2.0 to 2.5 H-bond; *purple*: 2.6 to 3. 5 H-bond; *green*: 3.6 and above; *blue*: hydrophobic. Note: this is an a.a. of CDR2 that is near the apex of the loop, in some cases closer contacts can be found at the bases of the loop; however, these contacts do not correlate with the location of the central cysteine; thus, relative position of the domain along the α-helix IP.

***Appendix for:** *Murray, J.S., 2023. Dichotomy in TCR V-domain dynamics binding the opposed inclined planes of MHC-II and MHC-I a-helices.*
